# Structures of TRPM5 channel elucidate mechanism of activation and inhibition

**DOI:** 10.1101/2021.03.25.437100

**Authors:** Zheng Ruan, Emery Haley, Ian J. Orozco, Mark Sabat, Richard Myers, Rebecca Roth, Juan Du, Wei Lü

**Author notes:** CORRESPONDING AUTHOR: Correspondence and requests for materials should be addressed to J. D., or W. L. These authors contributed equally to this work (Zheng Ruan, Emery Haley, Ian J. Orozco).

## Abstract

The Ca^2+^-activated TRPM5 channel plays an essential role in the perception of sweet, bitter, and umami stimuli in type II taste cells and in insulin secretion by pancreatic beta cells^1–3^. Interestingly, the voltage dependence of TRPM5 in taste bud cells depends on the intracellular Ca^2+^ concentration^4^, yet the mechanism remains elusive. Here we report cryo-electron microscopy structures of the zebrafish TRPM5 in an apo closed state, a Ca^2+^-bound open state, and an antagonist-bound inhibited state, at resolutions up to 2.3 Å. We defined two novel ligand binding sites: a Ca^2+^ binding site (Ca_ICD_) in the intracellular domain (ICD), and an antagonist binding site in the transmembrane domain (TMD) for a drug (NDNA) that regulates insulin and GLP-1 release^5^. The Ca_ICD_ site is unique to TRPM5 and has two roles: shifting the voltage dependence toward negative membrane potential, and promoting Ca^2+^ binding to the Ca_TMD_ site that is conserved throughout Ca^2+^-sensitive TRPM channels^6^. Replacing glutamate 337 in the Ca_ICD_ site with an alanine not only abolished Ca^2+^ binding to Ca_ICD_ but also reduced Ca^2+^ binding affinity to Ca_TMD_, suggesting a cooperativity between the two sites. We have defined mechanisms underlying channel activation and inhibition. Conformational changes initialized from both Ca^2+^ sites, 70 Å apart, are propagated to the ICD–TMD interface and cooperatively open the ion-conducting pore. The antagonist NDNA wedges into the space between the S1-S4 domain and pore domain, stabilizing the TMD in an apo-like closed state. Our results lay the foundation for understanding the voltage-dependent TRPM channels and developing new therapeutic agents to treat metabolic disorders.

## Introduction

Taste perception is one of the fundamental chemosensations in mammals, detecting the availability and the quality of food by converting the signal from tastants into electrical signals that the brain can interpret. Highly expressed in type II taste bud cells, the TRPM5 channel has been considered a key player in sensing sweet, umami, and bitter stimuli^1, 7^. TRPM5 is activated upon the elevation of cytoplasmic Ca^2+^ concentration that is caused by the binding of tastants to the taste receptors^8^. Activated TRPM5 then depolarizes the membrane and causes the CALHM1 channel to release the neurotransmitter ATP, which binds to the downstream P2X receptors that trigger the action potential of gustatory neurons, thus conveying taste information to the brain^8, 9^. TRPM5 also participates in other physiological processes in diverse cell types in a similar manner. For example, it is involved in insulin secretion by pancreatic beta cells^2, 3^ and in the immune response of tuft cells^10^. Thus, TRPM5 has broad implications for metabolic syndromes and immune disorders, and it is a potential drug target for the treatment of metabolic disorders such as obesity and type 2 diabetes^11^.

The transient <u>r</u>eceptor <u>p</u>otential superfamily, <u>m</u>elastatin subfamily (TRPM) consists of eight family members (TRPM1-8) that have diverse functional properties^12^. While most TRP family members are nonselective cation channels that are permeable to Na^+^ and Ca^2+^, TRPM5 and TRPM4 are the only two that are monovalent cation– selective and impermeable to Ca^2+^ (Ref ^13–15^). Moreover, TRPM5 and TRPM4 share substantial sequence similarity, and both are activated by intracellular Ca^2+^ in a voltage-and temperature-dependent manner; therefore, they have been classified as close homologs^16^. However, their biophysical properties vary with respect to Ca^2+^ sensitivity and ligand specificity, and the molecular basis for these differences is unknown^17^. For instance, TRPM5 is roughly 20-fold more sensitive to Ca^2+^ than TRPM4^17^. TRPM4 is inhibited by ATP, and its voltage dependence is modulated by decavanadate, while TRPM5 is insensitive to these ligands^17, 18^. By contrast, TRPM5, but not TRPM4, is modulated by the sweetener stevioside^19^. TRPM5 has also been an attractive pharmaceutical target for treating metabolic syndromes. For example, a family of small molecule TRPM5 inhibitors, including N’-(3,4-dimethoxybenzylidene)-2-(naphthalen-1-yl)acetohydrazide (NDNA), has been invented. These inhibitors have the potential to treat type II diabetes by enhancing insulin release and GLP-1 release^5^, which is contrary to the result of TRPM5 KO mice experiments^3^. Recently, structures of the voltage-dependent TRPM4 and TRPM8 channels have been reported^20–25^, but none have been captured in an active open state, preventing a detailed understanding of their gating mechanisms. To understand the molecular mechanisms by which Ca^2+^ activates and antagonist NDNA inhibits TRPM5, we performed electrophysiological and structural studies on TRPM5.

### Channel function, overall structures, and the ion-conducting pore

The zebrafish TRPM5 is highly sensitive to Ca^2+^, and 1 µM Ca^2+^ elicited a robust outward rectifying current in an excised inside-out patch (Extended Data Fig. 1a, k, u–w). Interestingly, the outward rectification of the TRPM5 currents apparently depends on the Ca^2+^ concentration. That is, at low Ca^2+^ concentration, the activation of TRPM5 requires membrane depolarization, whereas at high Ca^2+^ concentration, TRPM5 becomes markedly less voltage-dependent, having a nearly linear current–voltage relation (Fig. 1a; Extended Data Fig. 1a, k; Supplementary Fig. 1a). This unique property is conserved in human TRPM5 but not in its closest homologue TRPM4 (Extended Data Fig. 1b, h, l, r, u–w)^21^. Moreover, this shift in voltage dependence was previously observed with native TRPM5 currents in taste cells^4^. Our data imply that besides being an agonist, Ca^2+^ may also serve as a modulator to tune the voltage dependence of TRPM5.

**Figure 1:**
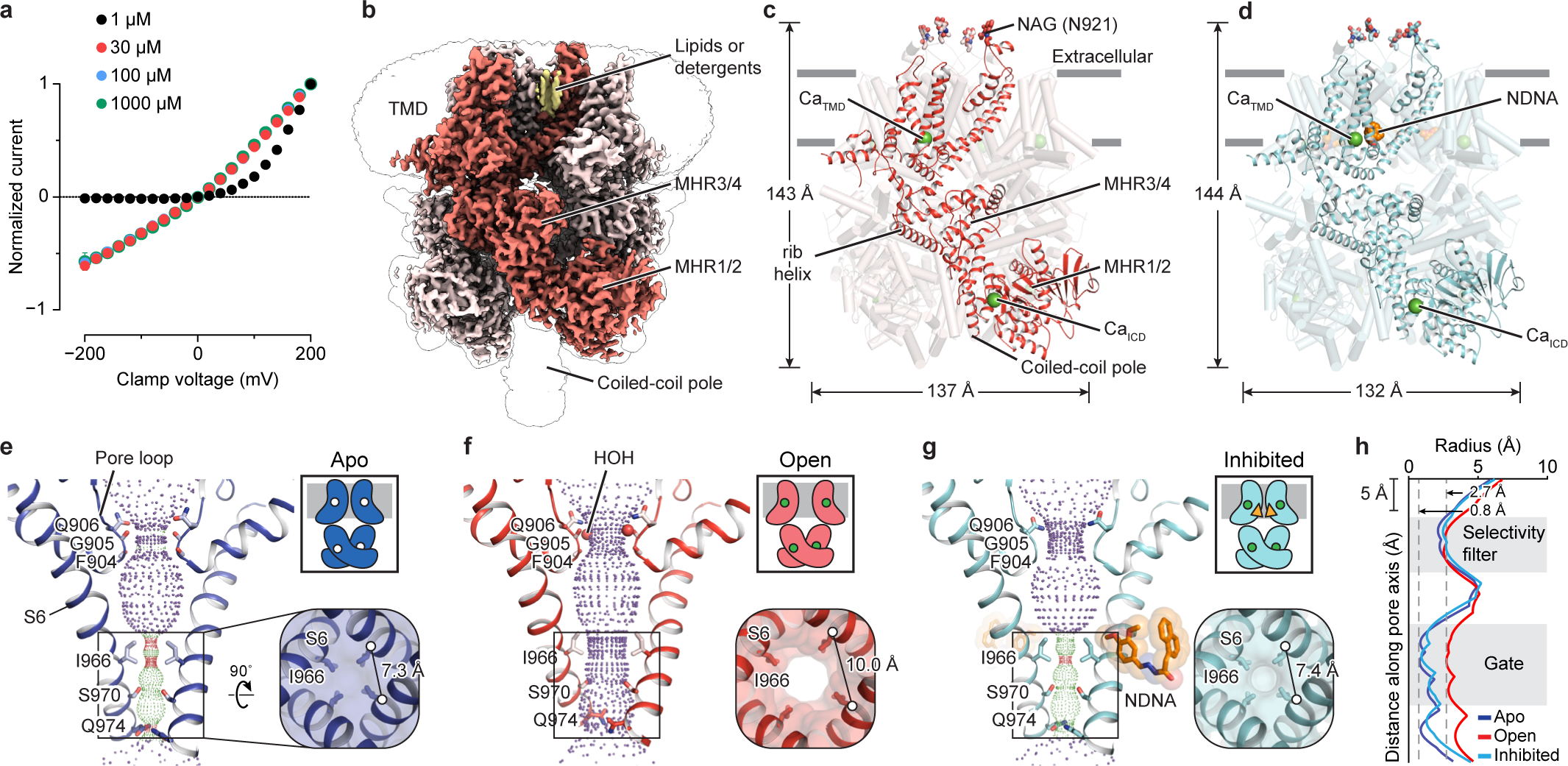
The overall architecture. **a**, Normalized Ca^2+^-activated currents from patches excised from tsA201 cells overexpressing zebrafish WT TRPM5 and recorded in the inside-out patch-clamp configuration. Bath solutions containing free Ca^2+^concentrations of 1, 30, 100, and 1000 µM were superfused and voltage clamps were imposed from +200 mV to –200 mV in steps of 20 mV with a final tail current pulse at –140mV.Background currents were subtracted by interleaved measurements with a calcium-free solution. Current amplitudes were measured at the end of the pulse (50 ms), normalized to +200 mV, and plotted as a function of clamp voltage (1 µM Ca^2+^ [*n* = 13 patches], 30 µM [*n* = 6], 100 µM [*n* = 4], 1000 µM [*n* = 3] from 12 transfections). See Extended Data Fig. 1a for representative traces. Tail current analysis was also performed (see Supplementary Figure 1). **b**, **c**, The cryo-EM map (**b**) and the atomic model (**c**) of Ca^2+^–TRPM5 viewed parallel to the membrane. One subunit is highlighted in red. The unsharpened reconstruction is shown as transparent envelope in (**b**). The Ca^2+^ ions are shown as green spheres in (**c**). **d**, The atomic model of NDNA/Ca^2+^–TRPM5 viewed parallel to the membrane. One subunit is highlight in cyan. The NDNA molecule is colored in orange. **e–g**, The profile of the ion-conducting pore in apo–TRPM5 (**e**), Ca^2+^–TRPM5 (**f**), and NDNA/Ca^2+^– TRPM5 (**g**) viewed parallel to the membrane. Purple, green, and red spheres define radii of >2.3, 1.2–2.3, and <1.2 Å, respectively. The pore region (shown in cartoon), residues (shown in sticks) forming the gate, and the selectivity filter in two subunits are depicted. Lower right panel: the channel gate viewed from the intracellular side; the distance between the Cα atoms of adjacent I966 residues is labeled. Upper right box: a cartoon representing two subunits of the apo state. The unoccupied/occupied Ca^2+^ sites are shown as unfilled/filled circles, respectively. The cell membrane is shown as gray. **h**, Plot of pore radius along the pore axis.

We solved the structures of zebrafish TRPM5 in both glyco-diosgenin (GDN) and lipid nanodiscs (Extended Data Figs. 2–5; Extended Table 1; Supplementary Fig. 2); the structures were virtually indistinguishable (Supplementary Fig. 2g). We focused on the TRPM5 in GDN because it produced cryo-EM maps of higher resolution. The apo, Ca^2+^-bound, and NDNA/Ca^2+^-bound structures were determined in the presence of 1 mM EDTA, 5 mM Ca^2+^ and 0.5 mM NDNA/5mM Ca^2+^ and had estimated resolutions of 2.9, 2.3, and 2.8 Å, respectively (Extended Data Figs. 3o, 4c, 5c). The maps were of excellent quality (Fig. 1b; Extended Data Fig. 6), which allowed us to *de novo* model nearly the entire protein (Fig. 1c-d), to identify two bound Ca^2+^ ions and a NDNA molecule in each subunit (Fig. 1c-d), two water molecules coordinating the Ca^2+^ in the transmembrane domain (TMD) and two water molecule in the pore loop region (Extended Data Fig. 7), and to unambiguously define the channel gate and the selectivity filter (Extended Data Fig. 7c-g).

The tetrameric TRPM5 is assembled with a TMD formed by six transmembrane helices and four characteristic intracellular melastatin homology regions (MHR1/2 and MHR3/4) (Extended Data Fig. 3e, f). Despite being the closest homologue to TRPM4, TRPM5 has a distinct monomeric structure, with the MHR1/2 domain tilting toward the TMD, resulting in a more compact tetrameric assembly and a different intersubunit interface (Fig. 1c; Extended Data Fig. 8a). Besides the conserved Ca^2+^ site in the TMD (Ca_TMD_), as observed in the structures of TRPM2, TRPM4, and TRPM8, we observed a novel Ca^2+^ binding site (Ca_ICD_) in the intracellular cytosolic domain at the interface between MHR1/2 and MHR3/4 domains (Fig. 1c).

The most remarkable difference between the apo and Ca^2+^-bound structures was at the ion-conducting pore (Fig. 1e, f, h). The intracellular half of the pore, which constitutes the channel gate, is restricted by I966 in the apo structure, giving a smallest radius of 0.8 Å (Fig. 1e, h); this represents an apo (resting) closed state (apo–TRPM5). By contrast, the Ca^2+^-bound structure has an enlarged pore with a smallest radius of 2.7 Å (Fig. 1f, h), which allows the passage of partially dehydrated monovalent cations, thus representing an agonist-bound active open state (Ca^2+^–TRPM5).

The extracellular half of the pore is confined by the pore loop, which shows little conformational change during channel gating and is generally considered to be responsible for ionic selectivity (Fig. 1e, f, h; Extended Data Fig. 8b). Here, we identified two ordered water molecules in each subunit (Extended Data Fig. 7c, e-f). Notably, the water molecules form a tight oxygen ring along with the backbone oxygen atoms of G905 (Fig. 1h), constituting the narrowest site in the pore loop region (Extended Data Fig. 7e, g). The radius is approximately 2.5 Å, about the Na^+^–O distance in a six-coordinate hydrated sodium (2.4 Å)^26^. We suggest this oxygen ring acts as the selectivity filter, and the four water molecules provide a favorable hydration layer for sodium ions to permeate^27^. A similar selectivity filter likely also exists in TRPM4, given by the conserved sequences (Extended Data Fig. 10) and structures of their pore loop (Extended Data Fig. 8b). The replacement of Q977 (equivalent to Q906 in zebrafish TRPM5) by a glutamate gives human TRPM4 a moderate permeability to Ca^2+^ (Ref ^28^). A glutamate at this position may attract Ca^2+^ ions by creating a binding site near the selectivity filter^29^. In addition, the glutamate might no longer form the same hydrogen bonding network, resulting in an altered conformation and size of the selectivity filter.

In the presence of NDNA and Ca^2+^ (NDNA/Ca^2+^–TRPM5), we observe a well-defined density in each subunit wedging into a cleft between the S1-S4 domain and the pore domain of the TMD (Fig. 1d). This density unambiguously hews to the shape of a NDNA molecule (Extended Data Fig. 9a, b). Of note, this binding site has not been reported for any of the TRPM family channels, thus representing a novel site for modulating channel activity. Despite Ca^2+^ binding to both Ca_TMD_ and Ca_ICD_, the TMD of NDNA/Ca^2+^–TRPM5 shows an apo-like conformation with a closed pore (Fig. 1d, g, h). We thus define it as an antagonist-bound inhibited state.

### Two calcium binding sites

The Ca_TMD_ site is located within the S1-S4 domain and is surrounded by four amino acids and two water molecules in an octahedral geometry (Fig. 2a). The key residues coordinating the Ca_TMD_ are absolutely conserved across the TRPM family members (Fig. 2c upper panel). Their replacement by an alanine largely abolished Ca^2+^-invoked currents (Extended Data Fig. 1x–z), which indicates that binding of Ca^2+^ to Ca_TMD_ is indispensable for the activation of zebrafish TRPM5. The integrity of Ca_TMD_ site is also important for the activity of rat TRPM5 and other Ca^2+^-dependent TRPM channels^30, 31^. We propose that the octahedral geometry of the Ca_TMD_ site is likely conserved among Ca^2+^-sensitive TRPM channels, in light of the high sequence conservation of the Ca_TMD_ site and the similar spatial organization of the Ca^2+^-coordinating residues (Fig. 2c; Extended Data Fig. 8c).

**Figure 2:**
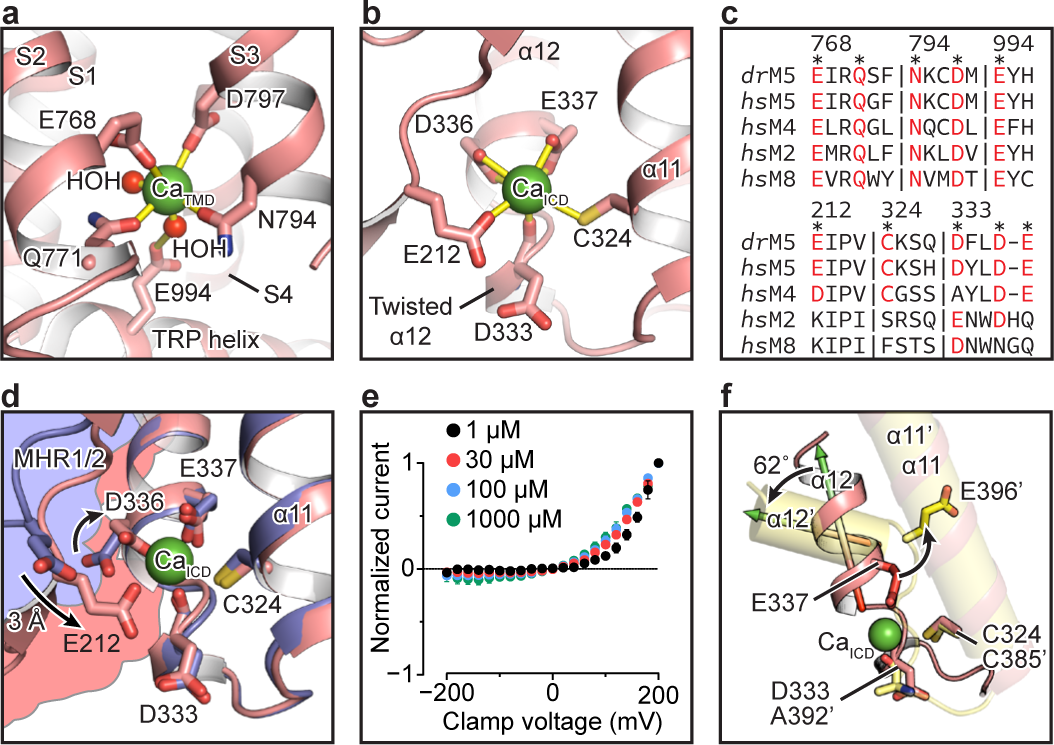
Ca^2+^ binding sites. **a**, **b**, The Ca_TMD_ (**a**) and Ca_ICD_ (**b**) sites. Ca^2+^ is shown as a green sphere. The coordinating residues and water molecules are shown in sticks and spheres, respectively. Polar interactions are indicated by yellow bars. **c**, Sequence alignment of the Ca_TMD_ (top) and Ca_ICD_ site (bottom) among zebrafish TRPM5 (*dr*M5), human TRPM5 (*hs*M5), human TRPM4 (*hs*M4), human TRPM2 (*hs*M2), and human TRPM8 (*hs*M8). The Ca^2+^ coordinating residues in zebrafish TRPM5 are indicated by asterisks, and conserved coordinating residues are in red. The residue numbers are according to zebrafish TRPM5 (UniProtID: S5UH55). Sequence segments are separated by vertical bars. **d**, Remodeling of the Ca_ICD_ site upon Ca^2+^ binding. Apo–TRPM5 and Ca^2+^–TRPM5 are in blue and red, respectively. Black arrows indicate the movement of E212 and D336. The MHR1/2 domains in apo–TRPM5 and Ca^2+^–TRPM5 are represented by blue and red surfaces, respectively, showing the movement of MHR1/2. **e**, Normalized current amplitudes from excised tsA cells overexpressing E337A mutant channels measured in the inside-out configuration as performed in Fig. 1a. The number of patches are (1 µM Ca^2+^ [*n* = 7 patches], 30 µM [*n* = 7], 100 µM [*n* = 4], 1000 µM [*n* = 4] from 8 transfections). See Extended Data Fig. 1c for representative traces. **f**, The superimposition of Ca_ICD_ site in TRPM5 (red) and TRPM4 (yellow, PDBID: 6BQR), by aligning the helix α11 and its equivalent in human TRPM4 (residues 396–403). The coordinating residues are shown as sticks. Equivalent structural elements and residues in TRPM4 are labeled with a prime symbol. The orientation of helix α12 and its equivalent (α12’) in human TRPM4 (residues 372–386) are indicated by colored 3D arrows. The differences between α11 and α11’, and between E337 and E396’, are indicated by black arrows.

The newly defined Ca_ICD_ site is at the interface between the MHR1/2 and MHR3/4 domains (Fig. 1c), in a negatively charged pocket. Part of the pocket is formed by a twisted helical segment, α12, of the MHR3 domain (Fig. 2b). The unique conformation of α12 allows the side chains of D336 and E337 on one side of the twist, and backbone oxygen of D333 on the other side, to face toward each other, thus creating a binding pocket to accommodate Ca^2+^ (Fig. 2b).

To investigate the mechanism underlying Ca_ICD_ binding, we carried out structural comparisons and electrophysiological experiments. The overlay of Ca^2+^–TRPM5 and apo– TRPM5 structures revealed that E337, C324, and the backbone oxygen of D333 form a rigid notch surrounding the binding site and show little movement upon Ca^2+^ binding; by contrast, D336 and E212 act as a flexible cap to enclose the Ca_ICD_ site (Fig. 2d). Specifically, E212 from MHR1/2 approaches the Ca_ICD_ site by moving approximately 3 Å, thus pulling the MHR1/2 domain closer to the MHR3 domain (Fig. 2d); meanwhile, the side-chain of D336 flips approximately 90° toward the Ca^2+^ (Fig. 2d). We propose that E337, the only negatively charged residue in the rigid notch, plays a key role in the binding of Ca^2+^. Indeed, replacement of E337 with an alanine (E337A) rendered TRPM5 voltage-sensitive even at high Ca^2+^ concentration (up to 1000 µM), distinct from the wild-type TRPM5, which becomes nearly voltage-independent at high Ca^2+^ concentration (Figs. 1b, 2e; Extended Data Fig. 1c, m, u–w; Supplementary Fig. 3b). Mutations of other coordinating residues to alanine only moderately altered the voltage sensitivity (Extended Data Fig. 1d–g, n–q, u–w; Supplementary Fig. 3c-f).

The Ca_ICD_ site is unique for TRPM5 because the residues coordinating Ca_ICD_ are conserved among TRPM5 orthologues, but not in other TRPM channels, except for TRPM4 (Fig. 2c lower panel). To understand why a site like the TRPM5 Ca_ICD_ was not observed in the published TRPM4 structures^20–23^ despite the conserved sequence, we compared their intracellular domains (ICDs) (Extended Data Fig. 10). We found that the key residues in TRPM4 are not close enough to each other to form a binding site because of two major structural differences. First, the structural element in TRPM4, which corresponds to the twisted helical segment α12 in TRPM5, is an intact α-helix, so that E396 (equivalent to E337 in TRPM5) cannot face the backbone oxygen of A392 (equivalent to D333 in TRPM5) (Fig. 2f). Second, the interface of MHR1/2 and MHR3, where the Ca_ICD_ site is located, has a markedly different arrangement than in TRPM5, manifested by different angles between helices α11 (on MHR1/2) and α12 (on MHR3) (Fig. 2f).

### Antagonist binding site

The antagonist NDNA is highly potent and inhibits Ca^2+^- induced TRPM5 currents with an IC_50_ of approximately 2.4 nM (Fig. 3a–c). The molecular structure of NDNA consists of two rings, a naphtalen and a dimethoxybenzylidene, which are linked by an acetohydrazide group (Extended Data Fig. 9a). In the NDNA/Ca^2+^–TRPM5 structure, the NDNA molecule is located at the interface between the S1-S4 domain and the pore domain (S5 and S6), near the Ca_TMD_ site (Fig. 3d). The two rings of NDNA are perpendicular to each other, forming a wedge shape. The naphtalen ring forms the base of the wedge, bracing on the S3 helix; the dimethoxybenzylidene ring forms the tip that penetrates through the cleft between S4 and S5, pressing against the pore domain of the adjacent subunit (Fig. 3d, e). The interaction between NDNA and TRPM5 is further enhanced by a hydrogen bond between the acetohydrazide linker and E853 on S5, and proximity of the same linker to W793 on S3 (Fig. 3f). Within the binding site, while most residues preserve their conformations in the apo state, the side chain of W869 flips to accommodate NDNA, forming a hydrogen bond with one of the methoxyl moieties on the dimethoxybenzylidene ring (Extended Data Fig. 9c; Fig. 3f).

**Figure 3:**
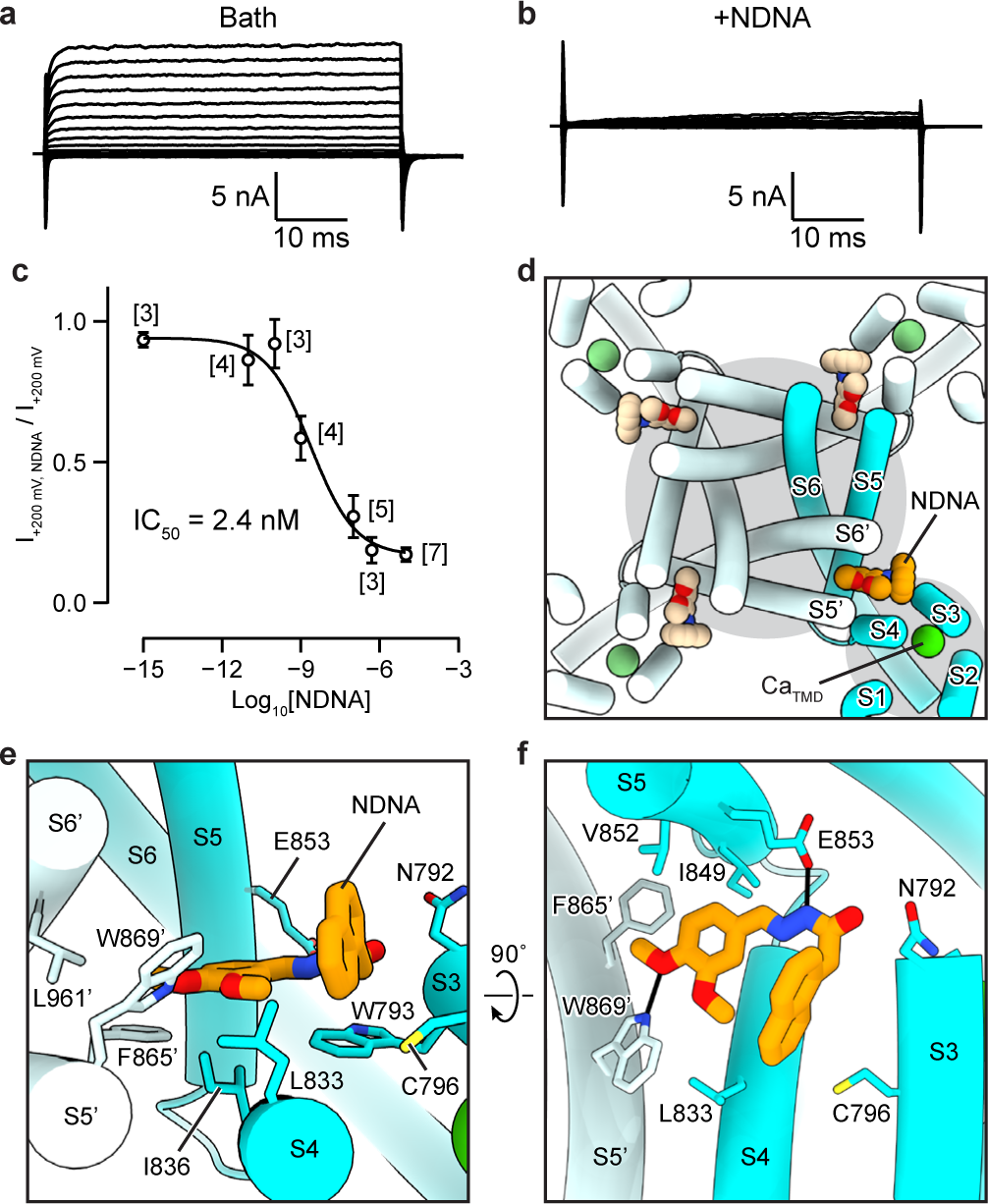
Effect and binding site of antagonist NDNA. **a** and **b**, Voltage-clamped (50 ms steps from +200 mV to -200 mV) calcium activated whole-cell currents from tsA201 cells over-expressing zebrafish WT TRPM5 were suppressed upon super-fusion of 10 µM NDNA in the bath solution. **c,** IC_50_ of NDNA, 2.4 nM, was determined by measuring and plotting the remaining current following inhibition (I_+200 mV_, _NDNA_ / I_+200 mV_, _bath_) using various NDNA concentrations (1 fM, 10 pM, 100 pM, 1 nM, 100 nM, 0.5 µM, 10 µM). Concentration is plotted in log (M). Each point represents the mean current, and bars indicate SEM. The number of cells is indicated in brackets. From non-linear fitting, the Hill Slope is -0.5, and the 95% CI is 0.5 – 23 nM. **d**, The pore domain of NDNA/Ca^2+^–TRPM5 viewed from the extracellular side. The four bound NDNA molecule is shown in orange. Transmembrane helices surrounding a copy of NDNA is labeled. Prime symbol indicates the adjacent subunit. Ca_TMD_ is shown in green sphere. **e** and **f**, Two close-up views for the detailed interactions mediated by the NDNA. One TRPM5 subunit is colored in cyan, whereas the adjacent subunit is colored in light cyan. Polar interactions between NDNA and residues are indicated by black lines.

In the NDNA/Ca^2+^–TRPM5 structure, although both Ca_TMD_ and Ca_ICD_ sites are occupied by Ca^2+^, we observed major differences relative to the Ca^2+^–TRPM5 open state. First, the ICD showed the same trend of motion relative to the apo–TRPM5 structure but to a lesser extent (Extended Data Fig. 9d). Second, the S1-S4 domain retained apo-like conformation and did not show marked conformational changes as observed in the Ca^2+^-bound open state, resulting in a different coordination of Ca_TMD_, with Q771 on the S2 helix not involved in the binding (Extended Data Fig. 9e, g). Lastly, the pore domain is closed, similar to the apo state (Extended Data Fig. 9h; Fig. 1g, h). Together, our data suggest that NDNA inhibits Ca^2+^-induced TRPM5 activation in a non-competitive manner. Despite Ca^2+^ binding, NDNA limits the movement of the S1-S4 domain and the pore domain, thus stabilizing the TMD in an apo-like closed state.

### The two roles of the Ca_ICD_ site

The alanine mutants of the key residues in the Ca_ICD_ site had the same Ca^2+^-induced channel activation as the wild type, but they shifted the voltage dependence toward a positive membrane potential to different degrees, with E337A having the strongest phenotype (Fig. 2e; Extended Data Fig. 1m–q, u; Supplementary Fig. 1). E337A remained voltage-dependent regardless of Ca^2+^ concentration, which was in sharp contrast to the wild type, in which current changed from voltage-dependent at low Ca^2+^ concentration to nearly voltage-independent at high Ca^2+^ concentration (Fig. 1b; Fig. 2e; Extended Data Fig. 1a, c, k, m, u–w; Supplementary Fig. 3a, b). We further looked into the same mutant on human TRPM5 (E351A) and observed the same phenotype (Extended Data Fig. 1h, i, r, s, u–w).

These results indicate that Ca_ICD_ modulates the voltage dependence of TRPM5. To ground our interpretations of the electrophysiological data, we determined the structure of E337A in the presence of 5 mM Ca^2+^ at 2.9 Å resolution (Fig. 4a; Extended Data Fig. 11). As expected, the ICD showed a wild type apo-like conformation and the Ca_ICD_ site was unoccupied (Fig. 4b, c). This supports the idea that replacement of E337 by an alanine indeed impaired Ca^2+^ binding to the Ca_ICD_ site and that the altered voltage dependence of E337A was caused by abolished Ca^2+^ binding to the Ca_ICD_ site.

**Figure 4:**
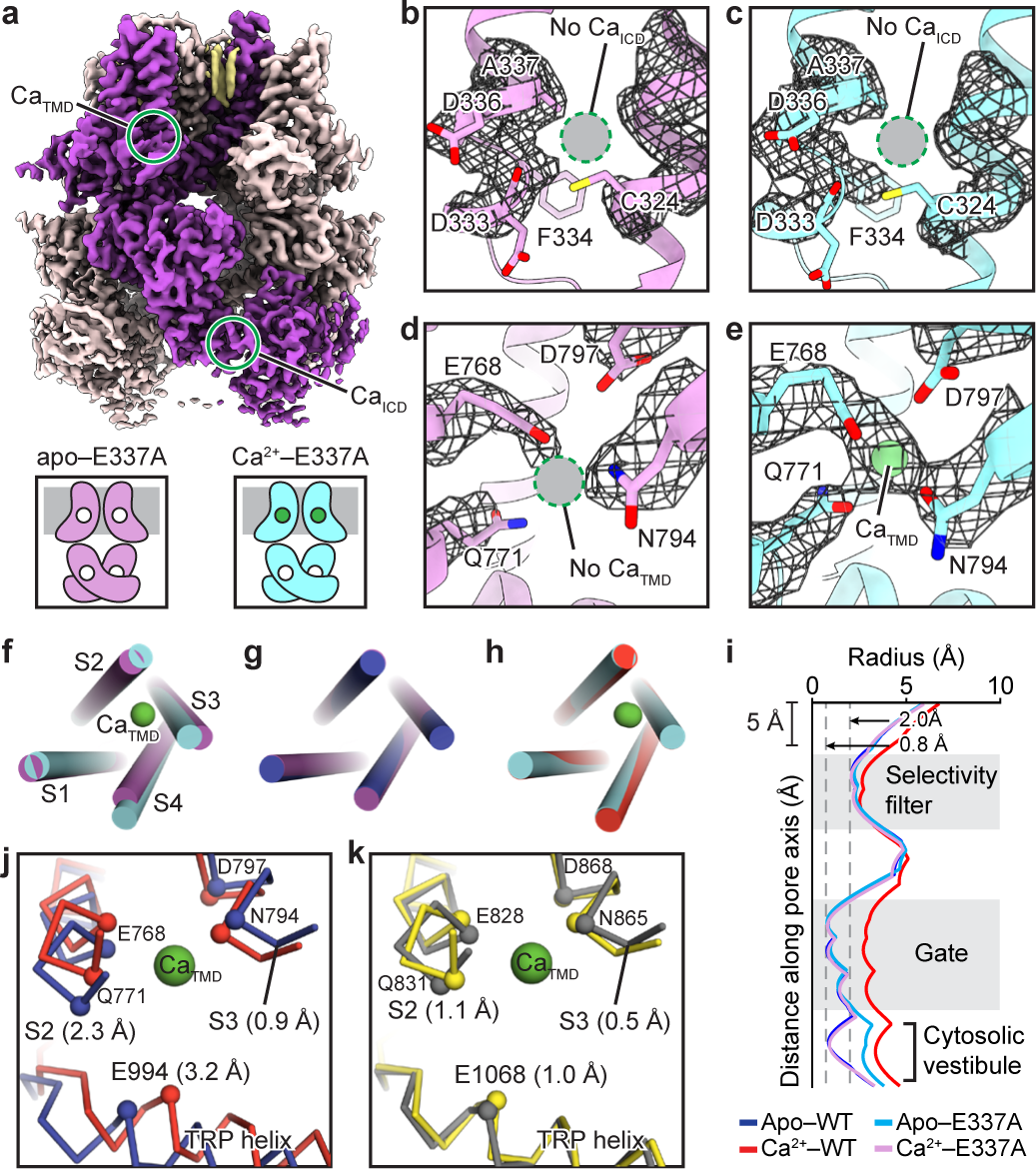
The structures of the Ca_ICD_-deficient mutant E337A. **a**, The upper panel shows the consensus map obtained from the Ca^2+^–TRPM5(E337A) data. One subunit is highlighted. The cartoons in the lower panel represent the two conformations that have distinct occupancies at the Ca_TMD_ site, obtained by single subunit analysis of the same data: apo– TRPM5(E337A) in magenta and Ca^2+^–TRPM5(E337A) in cyan. The unoccupied Ca^2+^ sites are shown as unfilled circles; occupied Ca^2+^ sites are shown as green circles. The cell membrane is represented in gray. **b**–**e**, the Ca_ICD_ site (**b**) and Ca_TMD_ site (**d**) in apo–TRPM5(E337A), and the Ca_ICD_ site (**c**), and Ca_TMD_ site (**e**) in Ca^2+^–TRPM5(E337A). The cryo-EM densities are shown in black mesh. The Ca^2+^ density is shown as a green sphere. Unoccupied sites are indicated by a dashed gray circle. **f**, The superimpositions of the S1-S4 domain of the apo–TRPM5(E337A) (magenta) and the Ca^2+^–TRPM5(E337A) (cyan) structures. **g**, The superimpositions of the S1-S4 domain of the apo–TRPM5(E337A) (magenta) and the apo–TRPM5 (blue) structures. **h**, The superimpositions of the S1-S4 domain of the Ca^2+^–TRPM5(E337A) (cyan) and the Ca^2+^–TRPM5 (red) structures. **i**, Plot of pore radius along the pore axis. **j**, **k**, Remodeling of the Ca_ICD_ site upon Ca^2+^ binding in TRPM5 (**j**) and TRPM4 (**k**). Apo–TRPM5 and Ca^2+^–TRPM5 are in blue and red, respectively (**j**). Apo–TRPM4 (PDBID: 6BQR) and Ca^2+^–TRPM4 (PDBID: 6BQV) are in gray and yellow, respectively (**k**). The Cα atoms of key residues and the Ca^2+^ are shown as spheres. Shown in parentheses are the distances of the root-mean-square-deviation (RMSD) between S2 (residues 767–772 in TRPM5 and 827–832 in TRPM4) and S3 (residues 793–798 in TRPM5 and 864–869 in TRPM4), and the distances of the Cα movements of E994 in TRPM5 and E1068 in TRPM4.

Although our data analysis workflow involved a focused classification of the TMD (Extended Data Fig. 11a), part of the TMD is still not unambiguously defined, and the densities for Ca_TMD_ were markedly weaker than those in the structure of Ca^2+^-bound wild-type TRPM5, indicating the structural heterogeneity of the TMD. We therefore performed structural analysis on a single subunit and obtained two conformations that had the same ICD but distinct S1-S4 domains in the TMD (Fig. 4b-h). One had an empty Ca_TMD_, termed apo–TRPM5(E337A) (Fig. 4b, d), and the other had an occupied Ca_TMD_, termed Ca^2+^–TRPM5(E337A) (Fig. 4c, e). This suggests that an unoccupied Ca_ICD_ site lowers the binding affinity of Ca^2+^ for the Ca_TMD_ site. The cooperativity between these two Ca^2+^ binding sites agrees with the observation that upon activation with 1 µM Ca^2+^, the current amplitudes of the E337A mutant at a clamp of +200 mV were substantially smaller (75%) than those of the wild type (Extended Data Fig. 1a, c, k, m, v).

Within the TMD, a closed ion-conducting pore was observed in the Ca^2+^–TRPM5(E337A) structure (Fig. 4i). This further supports the role of Ca_ICD_ as a voltage-modulating site, as its absence renders TRPM5(E337A) inactive due to the lack of membrane depolarization under the conditions for structural determination (Fig. 2e). Interestingly, the cytosolic vestibule, the part underneath the channel gate in Ca^2+^–TRPM5(E337A), is similar to that of Ca^2+^–TRPM5 (Fig. 4i), suggesting that the pore in Ca^2+^–TRPM5(E337A) may represent an intermediate state prior to channel opening.

The cooperativity between Ca_TMD_ and Ca_ICD_ sites implies that Ca_ICD_ might be physiologically relevant. To test this hypothesis, we determined the structure of wild-type TRPM5 in the presence of 6 µM Ca^2+^, a concentration similar to the Ca^2+^ EC_50_ of TRPM5 channels excised from native taste receptor cells (8 µM at –80 mV)^4^. Interestingly, this condition yielded both the apo conformation and the Ca^2+^-bound open conformation (Extended Data Fig. 4e-h). The ratio of protein particles belonging to the apo and open conformations, respectively, is 1.4:1. This data thus quantitatively correlates agonist-induced conformational changes to the EC_50_ determined by excised patch recordings. Furthermore, Ca_TMD_ and Ca_ICD_ are clearly occupied in the open state, which indicates that Ca^2+^ at EC_50_ concentration binds equally well to both sites, thus supporting the physiological relevance of Ca_ICD_ (Extended Data Fig. 4i).

To understand why the occupation of the Ca_ICD_ site is required for a high-affinity Ca_TMD_ site in TRPM5 but not in TRPM4, we compared the conformational changes in their Ca_TMD_ sites upon binding of Ca^2+^. In TRPM5, the helices S2 and S3—containing the coordinating residues and the TRP helix, which are key elements in transducing signals from the ICD to TMD— undergo substantial movement (Fig. 4j). By contrast, in TRPM4, the Ca_TMD_ site and the TRP helix mostly showed only minor sidechain rearrangement (Fig. 4k). This difference suggests that the Ca_TMD_ site in TRPM4 may be primed for Ca^2+^ binding, whereas in TRPM5, a high-affinity Ca_TMD_ site likely requires extensive rearrangement of the S2 and S3, assisted by Ca^2+^ binding to the Ca_ICD_ site.

Taken together, our data suggest the Ca_ICD_ site is physiologically relevant and has two important roles. First, it acts as a voltage modulator that shifts the voltage dependence toward negative potential, reminiscent of the effect of decavanadate and PIP2 on TRPM4^18, 32^. Second, it promotes Ca^2+^ binding to the Ca_TMD_ site and facilitates channel activation.

### Signal transduction from Ca_ICD_ to the TMD

The structural comparison of TRPM5 in the absence versus the presence of Ca^2+^ showed conformational rearrangement throughout the protein, with individual domains mostly showing rigid body movement (Fig. 5a). To understand the respective contributions of the two Ca^2+^ binding sites and how they cooperatively open the channel, we traced the conformational changes from the ICD and the S1-S4 domain to the ion-conducting pore.

**Figure 5:**
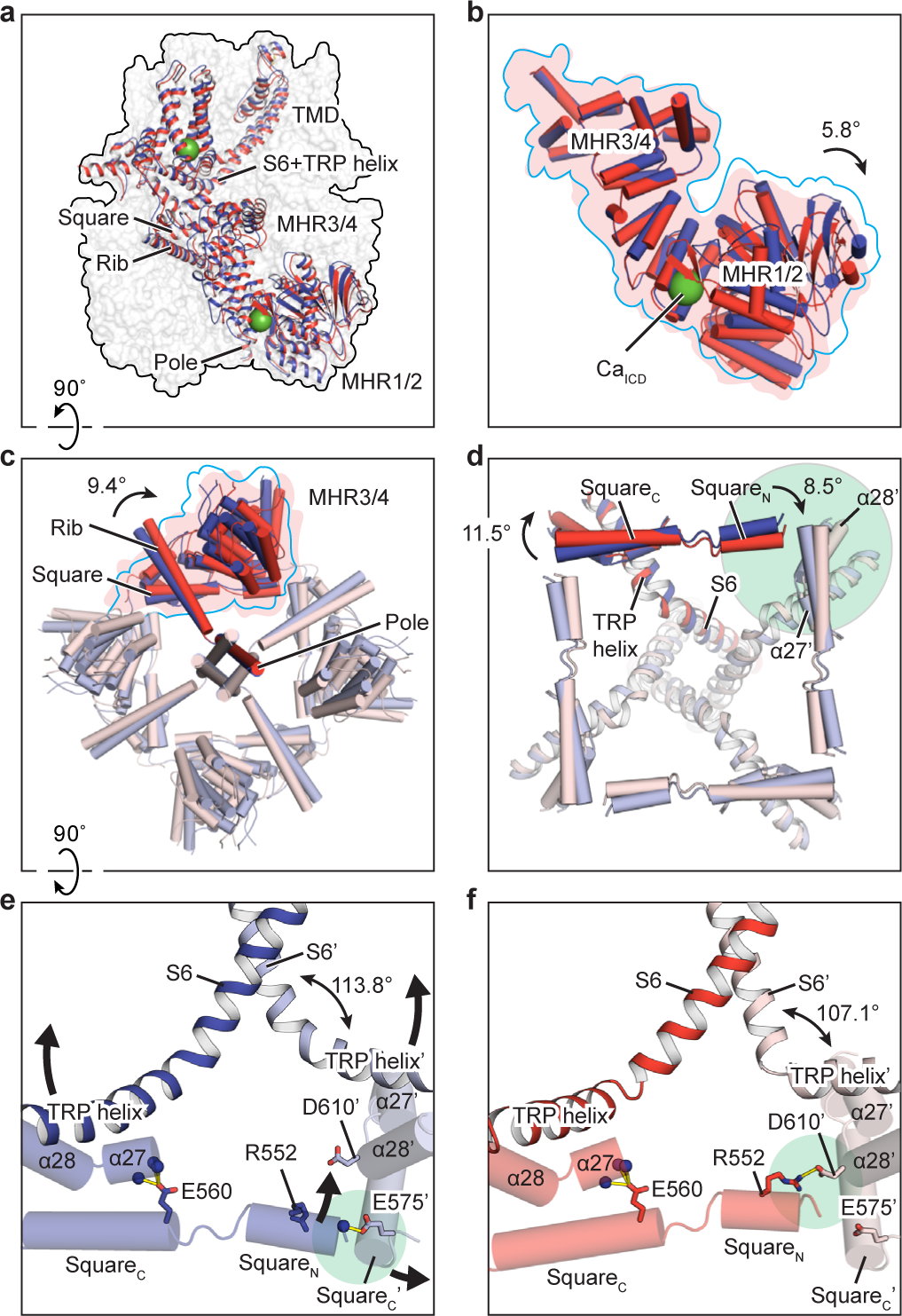
The signal transduction from ICD to TMD. **a**, The superimposition of apo– TRPM5 (blue) and Ca^2+^–TRPM5 (red) structures by aligning the coiled-coil poles in the C-terminal domain (CTD), viewed parallel to the membrane. The protein is shown in surface representation and one subunit is also shown in cartoon representation. Ca^2+^ is shown as green spheres. **b**, The superimposition of the MHR1-4 domains of apo–TRPM5 (blue) and Ca^2+^– TRPM5 (red) by aligning the MHR3/4 domain, viewed parallel to the membrane. The rotation of the MHR1/2 relative to the MHR3/4 domain upon Ca^2+^ binding is indicated. The surfaces are outlined in blue for apo–TRPM5 and filled with red for Ca^2+^–TRPM5. **c**, Superimposition of the MHR3/4 domain and the CTD rib and pole helices of apo–TRPM5 (blue) and Ca^2+^–TRPM5 (red) by aligning the CTD coiled-coil poles, viewed from the intracellular side. The surfaces of one subunit in both structures are shown in blue (apo–TRPM5) or in red (Ca^2+^–TRPM5). The rotation of the rib helices is indicated. **d**, The superimposition of the ICD–TMD interface of apo– TRPM5 (blue) and Ca^2+^–TRPM5 (red) by aligning the CTD coiled-coil poles (not shown), viewed from the intracellular side. The rotations of helices square_N_ and square_C_ are indicated. The green circle highlights the location of the intersubunit interface and ICD–TMD interface, as detailed in panels (**e**, **f**). **e**, **f**, The conformational rearrangement at the intersubunit interface and the ICD–TMD interface from apo–TRPM5 (**e**) to Ca^2+^–TRPM5 (**f**), viewed parallel to the membrane. Two adjacent subunits are shown in bright and light colors, respectively. Structural elements and residues of one subunit are labeled with a prime symbol. Interactions are shown in yellow bars. The single headed arrows indicate the movement of the square and TRP helices. The double headed arrow indicates the angle between S6 and TRP. The positions of the intersubunit interface are shown by green circles.

Clamped by two lobes, i.e., the MHR1/2 and MHR3/4 domains, the Ca_ICD_ site underwent an opening of 5.8° upon binding of Ca^2+^ (Fig. 5b). As a result, the rib helix, which penetrates into the interface between the MHR3/4 domain and the adjacent MHR1/2 domain, showed a clockwise rotation of 9.4° as viewed from the intracellular side (Fig. 5c). Meanwhile, just above the rib helix, four helices that form a square shape also rotated clockwise (Fig. 5d). Because these “square” helices mediate the contact between adjacent subunits, their rotation altered the intersubunit interface. Specifically, in the absence of Ca^2+^, the N- and C-termini of adjacent square helices are in close contact (Fig. 5e). Upon binding of Ca^2+^, this interface is disrupted as the adjacent termini rotate away from each other, resulting in a new interface between the N-terminus of the square helix and helix α28, where R552 and D610 form a salt bridge (Fig. 5f). Notably, the square helix in TRPM5 is broken into two short segments in the middle where E560 interacts with the N-terminus of helix α27 (Fig. 5e, f), a feature that is unique to TRPM5. By contrast, all the other TRPM channels have (or are predicted to have) a continuous square helix (Extended Data Fig. 8d, e).

We speculate that the square helix plays a role in the signal transduction from the Ca_ICD_ site to the TMD because it links the remodeling of the intersubunit interface upon Ca^2+^ binding to the TRP helix—a key element involved in channel gating—through helices α27 and α28. Indeed, replacement of E560 by an alanine, which presumably weakens the interaction between the square helix and helix α27, led to slower channel activation and deactivation kinetics (Extended Data Fig. 1j, t). Moreover, the R578Q polymorphism in human TRPM5 (which corresponds to position 561 on the square helix in zebrafish TRPM5) has been associated with obesity-related metabolic syndrome^33^.

### Channel opening by synergistic action of Ca_ICD_ and Ca_TMD_

Accompanying the conformational changes of the ICD induced by Ca^2+^ binding, the TRP helix is pushed toward the TMD where the Ca_TMD_ binding site is located (Fig. 6a). Here, surrounded by the TRP helix, S4-S5 linker, S2, and S3, is the region where the conformational changes induced by Ca^2+^ binding to the Ca_ICD_ and Ca_TMD_ sites meet, and where complex remodeling occurs (Fig. 6a, b). We performed a detailed structural analysis, and propose a mechanism by which the motion of the TRP helix promotes Ca^2+^ binding at the Ca_TMD_ site, ultimately leading to channel opening.

**Figure 6:**
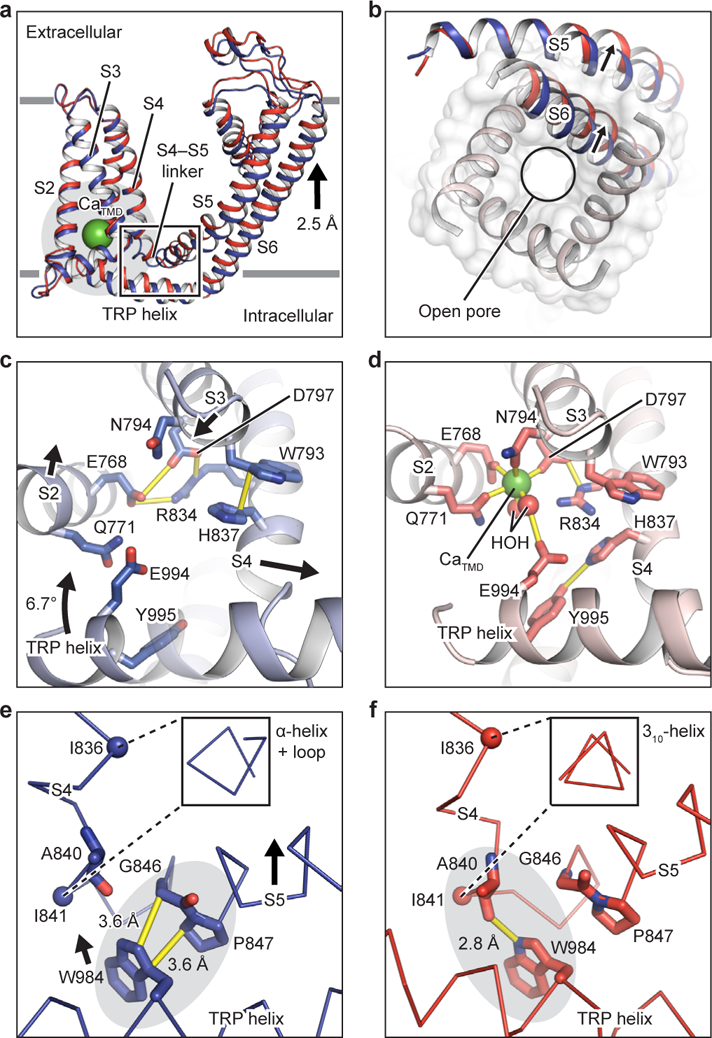
The channel opening. **a**, The superimposition of the TMD of a single subunit in apo–TRPM5 (blue) and Ca^2+^–TRPM5 (red) by aligning their S1-S4 domain, viewed parallel to the membrane. The center-of-mass movement of the pore domain is indicated. **b**, The superimposition of the pore domain in apo–TRPM5 (blue) and Ca^2+^–TRPM5 (red) by aligning their S1-S4 domain, viewed from the intracellular side. The pore domain of Ca^2+^–TRPM5 is shown in surface representation and the S5-S6 domain of one subunit from each structure is shown as a cartoon. The relative movements of helices S5 and S6 are indicated. **c**, **d**, Close-ups of the circled area in (**a**), viewed from the intracellular side. The remodeling of the Ca_TMD_ site from apo–TRPM5 (**c**) to Ca^2+^–TRPM5 (**d**). The movements of S2, S3, S4, and TRP helices are indicated by arrows. Interactions are shown in yellow bars. **e**, **f**, Close-ups of the boxed area in (**a**). W984 on the TRP helix switches its interaction partner from P847 and G846 in apo–TRPM5 (**e**) to I841 in Ca^2+^–TRPM5 (**f**). Interactions are shown in yellow bars. The movement of W984 is indicated. The contact area between the TRP helix and the S4-S5 linker is highlighted in grey. The segment between I836 and I841 turns into a 3_10_-helix in Ca^2+^–TRPM5. The inset shows the view along the axis of the S4 helix.

In the absence of Ca^2+^, i.e., when the Ca_ICD_ site is unoccupied, the Ca_TMD_ site is in a configuration that is difficult to access by Ca^2+^ because two crucial residues, E768 and D797, are locked by R834 in a triangular hydrogen bond network (Fig. 6c). This conformation is stabilized by the interaction between helices S3 and S4, with W793 and H837 stacking with each other (Fig. 6c). This explains why only a subset of particles from the Ca_ICD_ site-deficient mutant E337A showed an occupied Ca_TMD_ site even at high (5 mM) Ca^2+^ concentration (Extended Data Fig. 11a).

When the Ca_ICD_ site is occupied, the TRP helix tilts toward the TMD, leading to three consequences. First, it pushes the S4 helix away from S2 and S3 (Fig. 6c), allowing E768 and D797 to be readily released from R834 to coordinate Ca^2+^ together with Q771, N794, and two water molecules (Fig. 6d). Second, E994 on the TRP helix approaches the Ca_TMD_ site to coordinate one of the two water molecules, thus helping the Ca_TMD_ site bind Ca^2+^ (Fig. 6d). Third, Y995 on the TRP helix accommodates the flipped H837 by forming a hydrogen bond, thereby assisting in the decoupling of S4 from S3 by breaking the π-stacking between H837 and W793 (Fig. 6d). The decoupling of S4 from S3 is important, because it allows a relative movement between the S4-S5 linker and W984, a residue on the TRP helix that is absolutely conserved in the TRP superfamily and is crucial for channel gating^34–36^. As a result, the W984 switches its interaction partners from P847 and G846 on the N-terminus of S5 to the backbone oxygen of I841on S4, forcing the last turn of the S4 helix and part of the S4-S5 linker to stretch into a 3_10_-helix (Fig. 6e, f). The movement of W984 breaks the major interaction between the TRP helix and S5, eventually enabling the pore domain (helices S5 and S6) to relocate. Indeed, the structural comparison between the apo and open states of TRPM5 showed a movement of the pore domain by half an α-helical turn toward the extracellular side with an outward expansion, thus opening the ion-conducting pore (Fig. 6a, b).

### Conclusion and discussion

Our TRPM5 structures define two Ca^2+^ binding sites, Ca_ICD_ and Ca_TMD,_ which are 70 Å apart. Comparison of the structures in the apo and open states illustrates a molecular mechanism by which Ca^2+^ binding to the two locations synergistically leads to a complex conformational rearrangement at the interface between the ICD and the TMD (Fig. 7). This rearrangement eventually gives rise to the decoupling between the TRP helix and the S5 helix and the opening of the ion-conducting pore. We conclude that Ca_TMD_ functions as an orthosteric binding site for channel activation, while Ca_ICD_ modulates the voltage dependence and the accessibility of Ca_TMD_. The molecular basis by which the Ca_ICD_ site modulates the voltage dependence is still unclear; that will require the identification of the voltage sensor(s) and its working mechanism. We also defined a novel antagonist binding site in the TMD and elaborated a non-competitive inhibition mechanism by which the antagonist NDNA stabilizes the ion-conducting pore in an apo-like closed conformation (Fig. 7). Because NDNA is a potent TRPM5-selective antagonist and has potential implications in the treatment of diabetes, our work not only will facilitate the characterization of TRPM5 currents in many physiological processes, but is also important for the ongoing development of drugs targeting TRPM5.

**Figure 7:**
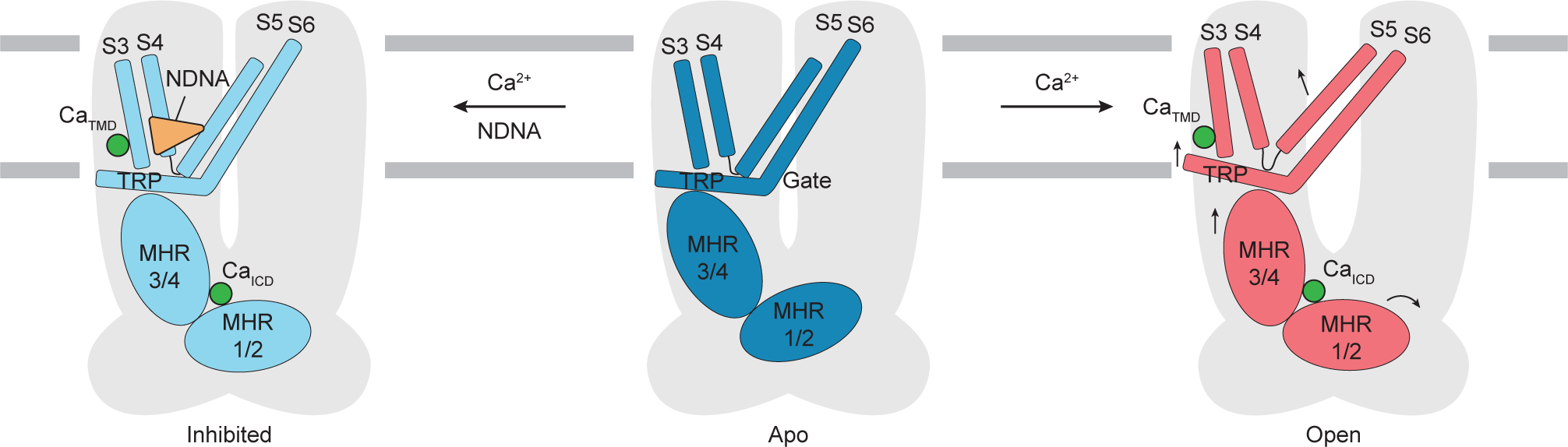
Schematic of the activation and inhibition mechanism of TRPM5. Conformational changes initialized from both Ca^2+^ sites cooperatively open the ion-conducting pore. The antagonist NDNA wedges into the space between the S1-S4 domain and pore domain, stabilizing the TMD in an apo-like closed state. The movements of individual structural elements are indicated by arrows.

TRPM4 and TRPM5 share substantial sequence similarity (55.7% between human TRPM4 and TRPM5, 58.1% between zebrafish TRPM4 and TRPM5), and both depolarize the cell membrane by sensing cytosolic Ca^2+^, but they are involved in different physiological processes. Interestingly, our data demonstrate that TRPM4 and TRPM5 are in fact structurally and functionally distinct. The unique Ca_ICD_ site endows TRPM5 with a complex gating and modulation mechanism by Ca^2+^, which may link to the physiological roles of TRPM5 in taste signaling and in the Ca^2+^ oscillation during insulin secretion by the pancreatic beta cells^2, 3^. We have elaborated a Ca^2+^-induced gating mechanism of a voltage-sensitive TRPM channel, which differs from that of the voltage-insensitive TRPM2^31, 37–39^. Our study highlights the important role of the ICD as a ligand-sensing domain in TRPM channels and lays a solid foundation for the development of novel therapeutic drugs that distinguish between TRPM4 and TRPM5.

### Data availability

The cryo-EM density map and coordinates of apo–TRPM5, Ca^2+^–TRPM5, NDNA/Ca^2+^– TRPM5, Ca^2+^–TRPM5(E337A) consensus, apo–TRPM5(E337A) single subunit and tetramer, Ca^2+^–TRPM5(E337A) single subunit and tetramer, apo–TRPM5(6μM Ca^2+^), Ca^2+^–TRPM5(6μM Ca^2+^), apo–TRPM5(nanodisc), and Ca^2+^–TRPM5(nanodisc) were deposited in the EMDB (Electron Microscopy Data Bank) under accession numbers EMD-xxxx, EMD-xxxx, EMD-xxxx, EMD-xxxx, EMD-xxxx, EMD-xxxx, EMD-xxxx, and EMD-xxxx. Atomic models for apo– TRPM5, Ca^2+^–TRPM5, NDNA/Ca^2+^–TRPM5, apo–TRPM5(E337A) tetramer, and Ca^2+^– TRPM5(E337A) tetramer were deposited in the Research Collaboratory for Structural Bioinformatics Protein Data Bank under accession codes xxxx, xxxx, xxxx, xxxx, and xxxx.

## Acknowledgements

We thank G. Zhao and X. Meng for the support with data collection at the David Van Andel Advanced Cryo-Electron Microscopy Suite. We appreciate the high-performance computing team of VAI for computational support. We thank D. Nadziejka and M. Martin for technical editing. W.L. is supported by National Institutes of Health (NIH) grants (R56HL144929, R01HL153219, and R01NS112363). J.D. is supported by a McKnight Scholar Award, a Klingenstein-Simon Scholar Award, a Sloan Research Fellowship in neuroscience, a Pew Scholar in the Biomedical Sciences award, and NIH grant (R01NS111031). Z.R. is supported by an American Heart Association postdoctoral fellowship (20POST35120556).

## Author Contributions

W.L. and J.D. supervised the project. E.H., Z.R., and B.R. generated TRPM5 mutants. Z.R. and E.H. carried out the purification, cryo-EM data collection, and processing. I.O. performed electrophysiological experiments. Z.R., J.D., and W.L. analyzed the structures. M.S. and R.M. synthesized the compound NDNA. Z.R., E.H., I.O., M.S., R.M., J.D., and W.L. contributed to the manuscript preparation. The authors declare no conflicts of interest.

## Methods

### TRPM5 expression and purification

Genes encoding full-length human and zebrafish TRPM5 (UniProtKB accession numbers Q9NZQ8, and S5UH5, respectively) were synthesized by Bio Basic and were sub-cloned into a pEG BacMam vector with an His8 tag, GFP, and a thrombin cleavage site at the N terminus^41^. Site-directed mutagenesis is performed by using QuikChange II Site-directed mutagenesis (Qiagen) or Q5 Site-Directed Mutagenesis (NEB) protocol, and confirmed via Sanger sequencing (Eurofins). For baculovirus production, each TRPM5 ortholog in a BacMam vector is transformed into DH10Bac cells, followed by P1 and P2 baculovirus generated in Sf9 cells. P2 viruses (8%) were used to infect tsA201 cells grown in Freestyle 293 Expression Medium in suspension culture (ThermoFisher). Infected cells were incubated for an initial 12 h at 37 °C before 10 mM sodium butyrate was added. Cells were then moved to a 30 °C incubator and allowed to grow for another 60 h with vigorous shaking. At 72 h post-infection, cells were harvested by centrifugation at 5000 rpm, 4 °C for 30 min. Cell pellets were washed with buffer containing 150 mM NaCl and 20 mM Tris pH 8.0 (TBS buffer) and stored at –80 °C.

Cell pellets from 200 ml culture were thawed on ice and resuspended in TBS buffer containing 1 mM PMSF, 0.8 μM aprotinin, 2 μg mL^−1^ leupeptin, 2 mM pepstatin A (which are all protease inhibitors) plus 1% GDN detergent (Anatrace). Protein was extracted from the membrane by whole-cell solubilization for 1 h at 4 °C with rotation. The solubilized protein was incubated with 2 mL TALON cobalt metal-affinity resin (Takara Bio) for 1 h. The TALON resin was then washed with 20 ml TBS buffer supplemented with 0.02% GDN and 15 mM imidazole. Protein was eluted with TBS buffer supplemented with 0.02% GDN and 250 mM imidazole. The eluent was concentrated to 500 μL and further purified by size-exclusion chromatography in TBS buffer containing 0.02% GDN. Peak fractions containing TRPM5 were pooled and concentrated to 5 mg/mL for grid freezing.

For nanodisc reconstitution, the eluent after immobilized metal affinity chromatography was mixed with MSP2N2 and soybean lipid extract at a molar ratio of 1:1:200 (TRPM5:MSP2N2:lipid). Three rounds of Bio-Beads (BIO-RAD) incubation at 4 °C was performed to facilitate nanodisc reconstitution. The Bio-Beads were then removed, and the sample was concentrated to 500 μL using an Amicon 100 kDa concentrator (MilliporeSigma). Size-exclusion chromatography was done in TBS buffer to further purify TRPM5-nanodisc complex. Peak fractions of TRPM5-nanodisc were collected and concentrated to 5 mg/mL for freezing grid.

### EM sample preparation and data acquisition

Freshly purified TRPM5 protein in GDN detergent was mixed with 1 mM EDTA (apo–TRPM5 and apo–TRPM5(E337A)), 5 mM Ca^2+^ (Ca^2+^–TRPM5 and Ca^2+^–TRPM5(E337A)) or 6 μM Ca^2+^ before grid preparation. For TRPM5-nanodisc sample, we added 0.05 mM digitonin to improve particle distribution on the grid. The apo–TRPM5(nanodisc) condition contains 1 mM EDTA, and the Ca^2+^–TRPM5(nanodisc) contains 1 mM Ca^2+^ and 0.5 mM steviol (Sigma). After mixing with the designated additives, a 2.5 μL aliquot of the sample was applied to a glow-discharged Quantifoil holey carbon grid (gold, 1.2/1.3 μm size/hole space, 300 mesh or gold, 2/1 μm size/hole space, 300 mesh), blotted for 1.5 s at 100% humidity using a Vitrobot Mark III, and then plunge-frozen in liquid ethane cooled by liquid nitrogen. The grids were loaded into a FEI Titan Krios transmission electron microscope operating at 300 kV with a nominal magnification of 130,000× and an energy filter (20 eV slit width). The apo–TRPM5, Ca^2+^–TRPM5(6 μM)(GDN), apo–TRPM5(nanodisc), and Ca^2+^–TRPM5(nanodisc) dataset was recorded by a Gatan K2 Summit direct electron detector in super-resolution mode with a binned pixel size of 0.521 Å. Each K2 movie was dose-fractionated to 40 frames for 8 s with a total dose of 49.6 e^−^ /Å^2^. The Ca^2+^–TRPM5, Ca^2+^–TRPM5(E337A), NDNA/Ca^2+^–TRPM5 datasets were collected by a K3 direct electron detector in super-resolution mode with a binned pixel size of 0.413 Å (K3). Each K3 movie was dose-fractionated to 75 frames for 1.5 s with a total dose of 47 e^−^/Å^2^. The automated image acquisition was facilitated using SerialEM^42^. The nominal defocus range was set from –0.9 μm to –1.9 μm.

### Cryo-EM data analysis procedure

The detailed workflow of data processing procedure is summarized in Extended Data Fig. 3-5, 11 and Supplementary Fig. 2. In general, the raw tif movie files for each dataset were motion-corrected and 2x binned using MotionCor2 v1.1.0 or RELION 3.0 (Ref ^43, 44^). The per-micrograph defocus values were estimated using Gctf 1.06 or ctffind 4.1 (Ref ^45, 46^). Particle picking was performed using gautomatch v0.56 (https://www2.mrc-lmb.cam.ac.uk/research/locally-developed-software/zhang-software/) or topaz v0.2.4 (Ref^47^) or topaz v0.2.4 (Ref ^48^). Junk particles were removed by 2D classification and heterogeneous refinement using CryoSPARC (v0.6 or v2.09). Selected good particles were then used to generate an initial 3D model by ab initio reconstruction followed by homogeneous refinement with C4 symmetry^49^. Multiple rounds of CTF refinement and Bayesian polishing were performed in relion to further improve the map resolution^50^.

At this stage, conformational heterogeneity was observed in the transmembrane domain (TMD) of the consensus refinement, indicating significant flexibility is present for TRPM5, especially in the extracellular pore loop area. To further improve the map quality, we performed focused classification by subtracting TMD signals from the particles^51^. After TMD-focused classification, we focused on the map with the highest nominal resolution and well-defined extracellular region for atomic model building.

For the Ca^2+^–TRPM5(E337A) dataset, conformational heterogeneity is still present in the transmembrane domain after TMD-focused classification. To overcome this issue, we performed symmetry expansion to the best particle set obtained from the TMD-focused classification and subtracted the single-subunit signals. Focused classification was conducted at the single-subunit level followed by 3D refinement. Two distinct conformations of the single subunit are identified for the Ca^2+^–TRPM5(E337A) dataset. The two conformation differ by the Ca^2+^ occupancy in the transmembrane domain, i.e., apo–TRPM5(E337A) and Ca^2+^–TRPM5(E337A). The single subunit maps were used for model building. To obtain tetrameric map for apo–TRPM5(E337A) and Ca^2+^–TRPM5(E337A), we further identified the homotetrameric TRPM5 particles that were solely composed of particles from each of the single subunit class. Although homo-tetrameric particles obtained after this procedure were very less, the refinement map still allowed us to generate a tetrameric model based on the single subunit map (see the model building section).

For the NDNA/Ca^2+^–TRPM5 dataset, conformational heterogeneity is observed in the ICD after TMD-focused classification. We then performed symmetry expansion (C4) and subtracted the ICD for each single subunit of TRPM5 particles. Subsequent 3D classification allowed us to obtain a homogeneous set of single TRPM5 subunit. We then identified homo-tetramer of TRPM5 that are consists of the homogeneous TRPM5 single subunit and refined the structure. This allowed us to obtain a map with better defined ICD to assist model building, despite with slightly worse nominal resolution compared to the consensus refinement before ICD classification.

For all dataset, the Gold standard Fourier shell correlation (FSC) 0.143 criteria were used to provide the map resolution estimate^52^. The cryo-EM maps were visualized using UCSF ChimeraX^53^.

### Model building

The atomic model for apo–TRPM5 was built into the cryo-EM density manually using Coot v0.89 and subjected to real-space refinement in Phenix^54, 55^. The apo–TRPM5 model contained residues 16-429, 446-473, 489-653, 698-1020, and 1027-1092. One GDN molecule lacking one of the two maltose groups (GDP), and one diosgenin molecule (DIO) were modeled into the lipid- or detergent-like densities for each chain. The geometrical restraints for DIO, GDP and NDNA were generated using the Grade Web Server (http://grade.globalphasing.org). Glycosylation at N921 was modeled as N-acetyl-beta-D-glucosamine (NAG). The Ca^2+^–TRPM5 open models were built by first docking the apo-TRPM5 closed model into the corresponding cryo-EM map density and adjusted manually in Coot. Two Ca^2+^ atoms were added to the TMD and ICD Ca^2+^ binding sites of the Ca^2+^–TRPM5 model. The Ca^2+^–TRPM5 open model was of sufficient resolution to allow us further place two ordered water molecules in the TMD Ca^2+^ binding site, and two water molecules in the selectivity filter for each chain. The apo– TRPM5(E337A) and Ca^2+^–TRPM5(E337A) model were first built based on the cryo-EM maps of the single-subunit refinement result. The single-subunit model was then rigid-body-fitted into the tetramer cryo-EM maps reconstructed from homo-tetrameric TRPM5 particles. We did not build atomic model for Apo–TRPM5(nanodisc) and Ca^2+^–TRPM5(nanodisc) dataset because these maps are identical to the corresponding maps from the GDN detergent conditions (Extended Data Fig. 5g). It is worth mentioning that although we included 0.5 mM steviol when preparing the Ca^2+^–TRPM5(nanodisc) grid, we are not able to identify density that corresponds to the steviol molecule.

### Electrophysiology

In the inside-out patch clamp configuration, voltage-clamped membrane currents were measured from tsA201 cells overexpressing plasmids encoding N-terminal GFP tagged WT and mutant TRPM5 channels from zebrafish and human. Following 1 d post-transfection with Lipofectamine 2000, cells were trypsinized and replated onto poly-L-lysine-coated (Sigma) glass coverslips. After cell adherence, the coverslips were transferred to a low-volume recording chamber with a pH 7.4 bath solution containing (in mM) 150 NaCl, 3 KCl, 10 HEPES, 2 CaCl_2_, 1 MgCl_2_, and 12 mannitol. Cells with fluorescence at the plasma membrane were patched with pipettes containing a pH 7.4 solution of (in mM) 150 NaCl, 10 HEPES, and 5 EGTA. Upon tight-seal formation, the bath solution was super-fused with the calcium-free EGTA solution. Following excision, patches were exposed using a manifold to super-fused bath solutions containing various free calcium concentrations. For preparing 1, 20, 100, or 1000 μM of free calcium, 4.46, 5.01, 5.1, or 6 mM CaCl_2_ was added to a pH 7.4 solution of 150 mM NaCl, 10 mM HEPES, 5 mM EGTA. Free calcium concentrations were calculated with https://somapp.ucdmc.ucdavis.edu/pharmacology/bers/maxchelator/CaEGTA-TS.htm. At room temperature (21-23 °C), patches from a holding voltage of 0 mV were clamped (Clampex 11.0.3, Multiclamp 700 B) using 50-ms steps from +200 mV to -200 mV (intracellular side relative to extracellular) with a final tail pulse at -140 mV. Electrical signals were digitized at 10 kHz and filtered at 2 kHz. Typically, measurements of TRPM5 activation by individual bath solutions containing various calcium concentration were interleaved with measurements where the bath solution was superfused with calcium-free EGTA solution. Using offline analysis (ClampFit 11.0.3) the currents in the absence of calcium were then subtracted from currents measured in the presence of calcium to acquire specific calcium-activated currents. Current amplitudes were measured at the end of the pulse. For normalizing current, the clamp at + 200 mV was chosen. Whole-cell measurements were performed in tsA201 cells following 1d transfection of zebrafish TRPM5 Ca_TMD_ mutant channels. Patch pipettes were filled with a 1 μM free calcium concentration solution (pH 7.4) composed of (in mM): 150 NaCl, 1 MgCl_2_, 10 Hepes, 5 EGTA, 4.45 CaCl_2_. The bath solution (pH 7.4) contained (in mM) 150 NaCl, 3 KCl, 10 Hepes, 2 CaCl_2_, 1 MgCl_2_, and 12 mannitol. Voltage clamps (50ms steps from +200 to -200 mV) were imposed approximately 1 minute after the whole cell configuration was acquired. Analysis was performed with GraphPad.

For determining the IC50 of NDNA, whole cell current analysis was performed where TRPM5 currents were evoked with 1 µM calcium in the patch pipette (as described above). Upon whole cell acquisition, currents were first measured in bath solution and then re-measured 30-60 s following super-fusion of bath solution containing various NDNA concentrations (1 fM, 10 pM, 100 pM, 1 nM, 100 nM, 0.5 µM, 10 µM). NDNA was stored at 50 mM (DMSO) and serially diluted using bath solution. For each cell measured, only one concentration of NDNA was tested. Inhibition kinetics was monitored using a step protocol (+100 mV) and typically complete (steady-state current) within a minute. Inhibited current was plotted as a function of NDNA concentration and fitted using Prism software (inhibitor versus response, variable slope equation).

### Preparation of *N*’-(3,4-dimethoxybenzylidene)-2-(naphthalen-1-yl)acetohydrazide (compound NDNA)

N’-(3,4-dimethoxybenzylidene)-2-(naphthalen-1-yl)acetohydrazide (NDNA) was synthesized according to the US Patent US8193168 (Bryant et al., 2008) (Supplementary Figure 4a). Briefly, A solution of commercial available ethyl 2-(naphthalen-1-yl)acetate (compound 1) (20 g, 93.34 mmol, 1 *eq*), NH_2_NH_2_.H_2_O (9.54 g, 186.69 mmol, 9.26 mL, 2 *eq*) in EtOH (100 mL) was stirred at 80°C for 16 h. TLC (Petroleum ether/Ethyl acetate = 2/1) showed that most of the compound 1 (R_f_ = 0.5) was consumed and a new spot (R_f_ = 0.05) was given. The reaction mixture was concentrated under vacuum to give white solid. The white solid was triturated with Petroleum ether/Ethyl acetate = 4:1 (100 mL) for 10 min. The mixture was filtered, and the filter cake was dried under vacuum to give the tittle compound 2 (12.8 g, 59.58 mmol, 63.8% yield, 93.2% purity) as a white solid. ^1^H NMR (400 MHz, DMSO-*d*_6_, see Supplementary Figure 4b upper panel) δ ppm 9.34 (s, 1H), 8.13 (d, *J* = 8.4 Hz, 1H), 7.92 (d, *J* = 7.6 Hz, 1H), 7.81 (d, *J* = 5.6 Hz, 1H), 7.54-7.51 (m, 2H), 7.45-7.44 (m, 2H), 4.25 (s, 2H), 3.84 (s, 2H). LCMS 0-60% ACN–H_2_O, ESI + APCI: Rt = 0.767 min, m/z = 201.1 (M+H)^+^.

A solution of 2-(naphthalen-1-yl)acetohydrazide (compound 2) (10.8 g, 50.27 mmol, 1 *eq*) and 3,4-dimethoxybenzaldehyde (compound **3**) (8.35 g, 50.27 mmol, 1 *eq*) in EtOH (50 mL) was stirred at 80°C for 30 min. A white solid separated out. TLC (Petroleum ether /Ethyl acetate = 1/1) showed that compound 2 (R_f_ = 0.1) was consumed and a major spot (R_f_ = 0.3) formed. The reaction mixture was cooled to 20°C. 100 mL EtOH was added to above solution and stirred for 10 min. The mixture was filtered and the filter cake was dried under vacuum to give NDNA (9.20 g, 26.33 mmol, 52.3% yield, 99.7% purity) as a white solid. ^1^H NMR (400 MHz, DMSO- *d*_6_, see Supplementary Figure 4b lower panel) δ ppm 11.06-11.38 (m, 1H), 8.18 (m, 1.4H), 7.96 (m, 1.6H), 7.93 (m, 1H), 7.55-7.48 (m, 4H), 7.31-7.30 (m, 1H), 7.18 (m, 1H), 7.01-6.99 (m, 1H), 4.53-4.23 (m, 2H), 3.80-3.65 (m, 6H). LCMS 5-95% ACN–H_2_O, ESI + APCI Rt = 0.826 min, m/z = 349.0 (M+H)^+^.

## Extended Data Figure legends

**Extended Data Figure 1:**
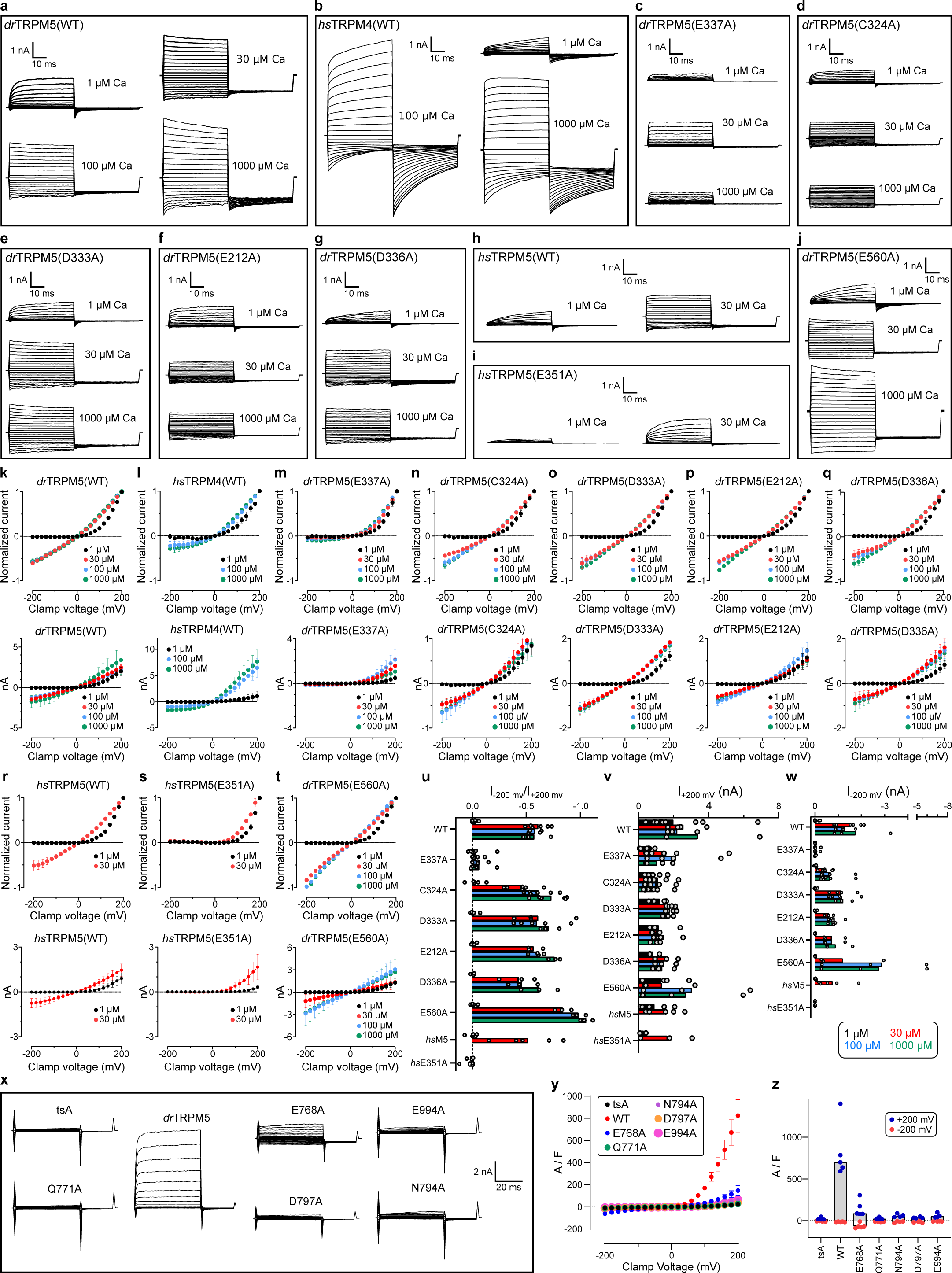
Patch-clamp analysis of TRPM5 and TRPM4 channels. Representative current traces of inside-out patch-clamp measurements from tsA201 cells overexpressing human TRPM4 (*hs*TRPM4), human TRPM5 (*hs*TRPM5), and zebrafish TRPM5 (*dr*TRPM5) channels. Patches were stimulated with either 1, 30, 100, or 1000 μM Ca^2+^ and were voltage-clamped from +200 mV to –200 mV. See Methods for detailed description. The number of patches and transfections were **a,** *dr*TRPM5(WT) 1 µM Ca^2+^ [*n* = 13 patches], 30 µM [6], 100 µM [4], 1000 µM [3] from 12 transfections; **b,** *hs*TRPM4(WT): 1 µM Ca^2+^ [4], 100 µM [4], 1000 µM [4] from 3 transfections; **c,** *dr*TRPM5(E337A): 1 µM Ca^2+^ [6], 30 µM [6], 100 µM [4], 1000 µM [4] from 8 transfections; **d,** *dr*TRPM5(C324A): 1 µM Ca^2+^ [6], 30 µM [6], 100 µM [6], 1000 µM [6] from 4 transfections; **e,** *dr*TRPM5(D333A): 1 µM Ca^2+^ [5], 30 µM [4], 100 µM [5], 1000 µM [3] from 3 transfections; **f,** *dr*TRPM5(E212A): 1 µM Ca^2+^ [4], 30 µM [3], 100 µM [4], 1000 µM [3] from 3 transfections; **g,** *dr*TRPM5(D336A): 1 µM Ca^2+^ [3], 30 µM [3], 100 µM [3], 1000 µM [3] from 2 transfections; **h,** *hs*TRPM5(WT): 1 µM Ca^2+^ [5], 30 µM [5] from 3 transfections; **i,** *hs*TRPM5(E351A): 1 µM Ca^2+^ [3], 30 µM [3] from 1 transfection; and **j,** *hs*TRPM5(E560A): 1 µM Ca^2+^ [5], 30 µM [3], 100 µM [3], 1000 µM [3] from 5 transfections. Currents measured in the absence of calcium were subtracted from currents measured in the presence of various calcium concentrations. **k**–**t**, Mean current amplitudes (50 ms) of experiments (**a**–**j**) were plotted as a function of clamp voltage. The +200 mV clamp was chosen for normalization. Horizontal bars represent SEM. In some cases the symbol size is larger than the error bars. The normalized current-voltage relation plots of *dr*TRPM5(WT) and *dr*TRPM5(E337A) are identical to those presented in Fig. 1a and Fig. 2e, respectively. **u**–**w**, Individual patch clamp measurements I_–200mV_ / I_+200mV_, I_+200mV_, I_–200mV_, of experiments (**a**–**j**) are shown as individual points, where bars represent mean values. **x**, Representative whole-cell current traces of tsA overexpressing WT and Ca_TMD_ mutant *dr*TRPM5 channels. Clamps were imposed from +200 mV to –200 mV. The number of cells measured were tsA201 [*n* = 4 cells], *dr*TRPM5(WT) [5], *dr*TRPM5(E768A) [5], *dr*TRPM5(Q771A) [4], *dr*TRPM5(N794A) [4], *dr*TRPM5(D797A) [4], and *dr*TRPM5(E994A) [4] from 2–3 transfections. **y**, Mean current amplitudes of experiments in (**x**) were measured at 50 ms and plotted as a function of clamp voltage. Horizontal bars represent SEM. **z**, Individual measurements at clamps of +200 mV (I_+200mV_) and -200 mV (I_–200mV_) of experiments in (**x**) are shown as individual points, with bars representing mean values.

**Extended Data Figure 2:**
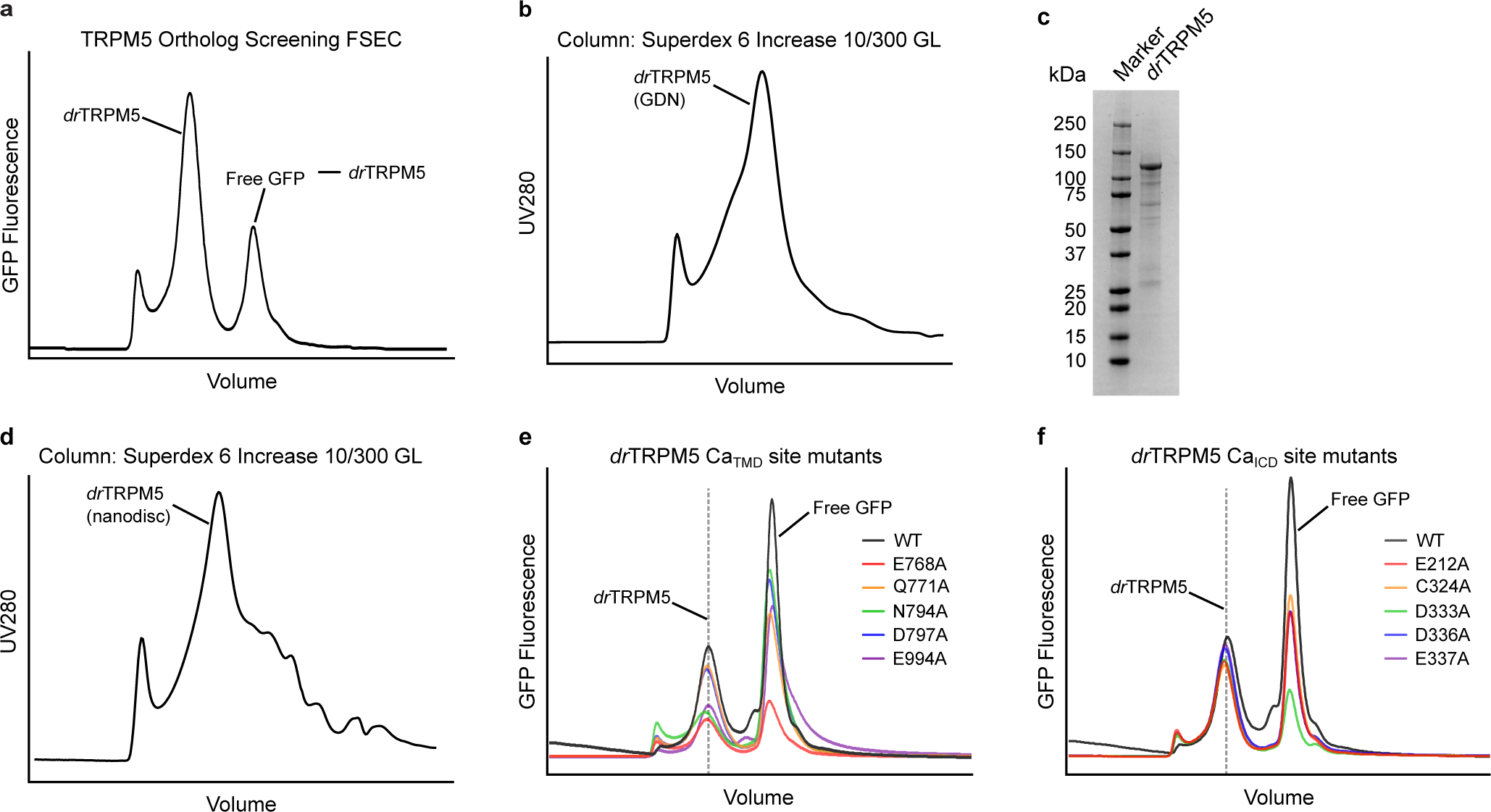
TRPM5 detergent screening, purification, and expression test of calcium-binding-site mutants. **a**, Fluorescence size-exclusion chromatography (FSEC) analysis of GFP-tagged zebrafish TRPM5 (*dr*TRPM5). The whole-cell sample was solubilized using GDN detergent and injected into a Superdex 6 Increase 5/150 GL column for high-performance liquid chromatography (HPLC) analysis. GFP fluorescence signal was probed across the retention volume. **b**, The size-exclusion chromatography profile of purified *dr*TRPM5 in GDN using Superdex 6 Increase 10/300 GL column. **c**, The SDS gel of purified *dr*TRPM5 protein. The uncropped raw gel image is provided in Supplementary Fig. 1. **d**, The size-exclusion chromatography profile of *dr*TRPM5 reconstituted into lipid nanodiscs. **e** and **f**, The FSEC profiles of the *dr*TRPM5 Ca_TMD_ and Ca_ICD_ site mutants, respectively. The expected retention volume for the *dr*TRPM5 protein is indicated.

**Extended Data Figure 3:**
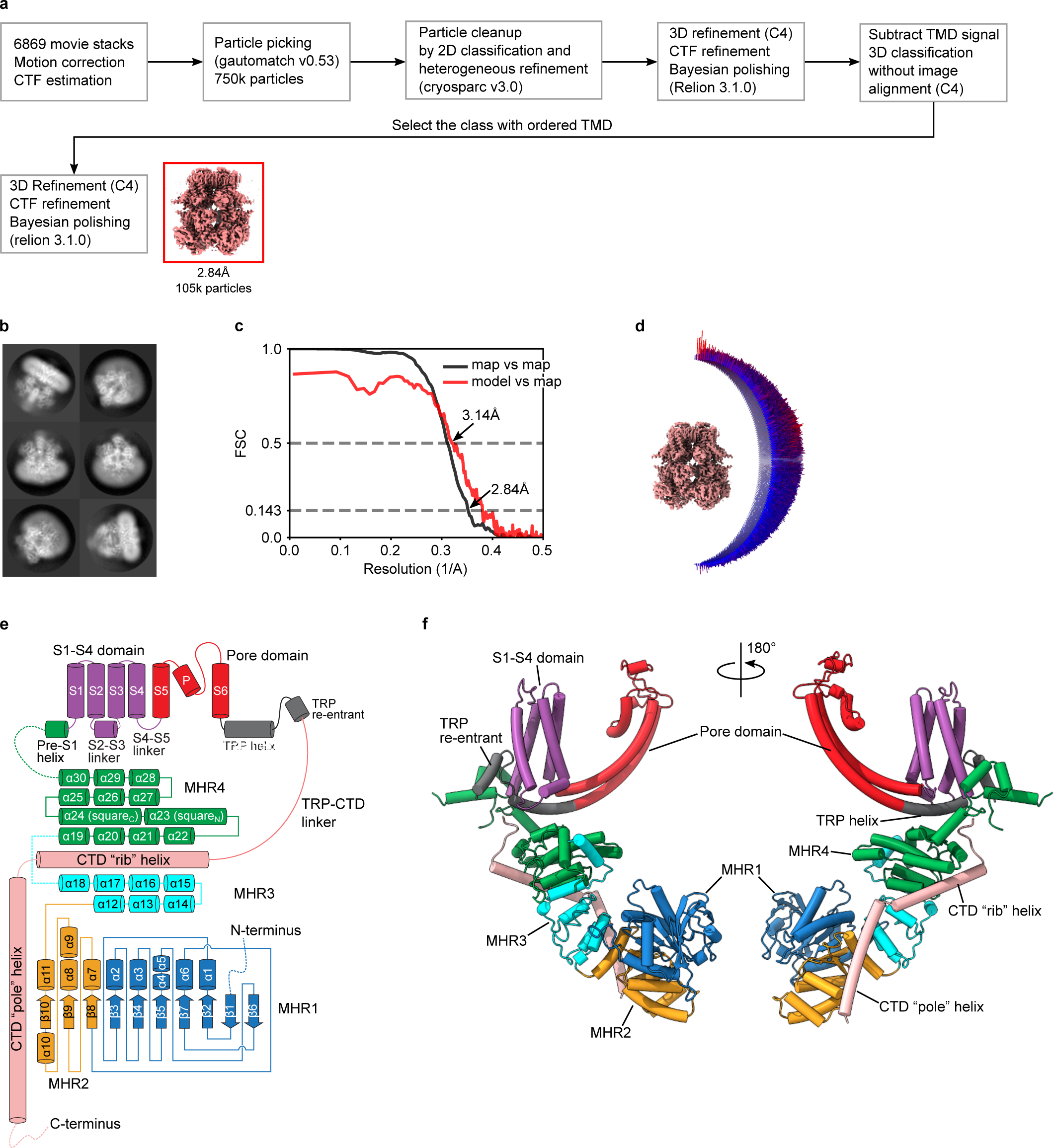
Apo–TRPM5 in GDN detergent. **a**, The data processing workflow for apo–TRPM5 dataset. **b**, The representative 2D class average of apo–TRPM5. **c**, The Fourier shell correlation (FSC) curves for the apo–TRPM5. The cryo-EM map FSC is shown in black and the model vs. map cross-correlation is shown in red. The map resolution was determined by the gold-standard FSC at 0.143 criterion, whereas the model vs. map resolution was determined by a correlation threshold of 0.5. **d**, The angular distribution of particles that gave rise to the apo–TRPM5 cryo-EM map reconstruction. **e**, A schematic domain organization of a single TRPM5 subunit. Secondary structures and important domains are labeled. **f**, The atomic model of a single TRPM5 subunit in cartoon representation. The domains are colored as in (**e**). The left and right panels are two different views of the same subunit rotated 180° along the central axis.

**Extended Data Figure 4:**
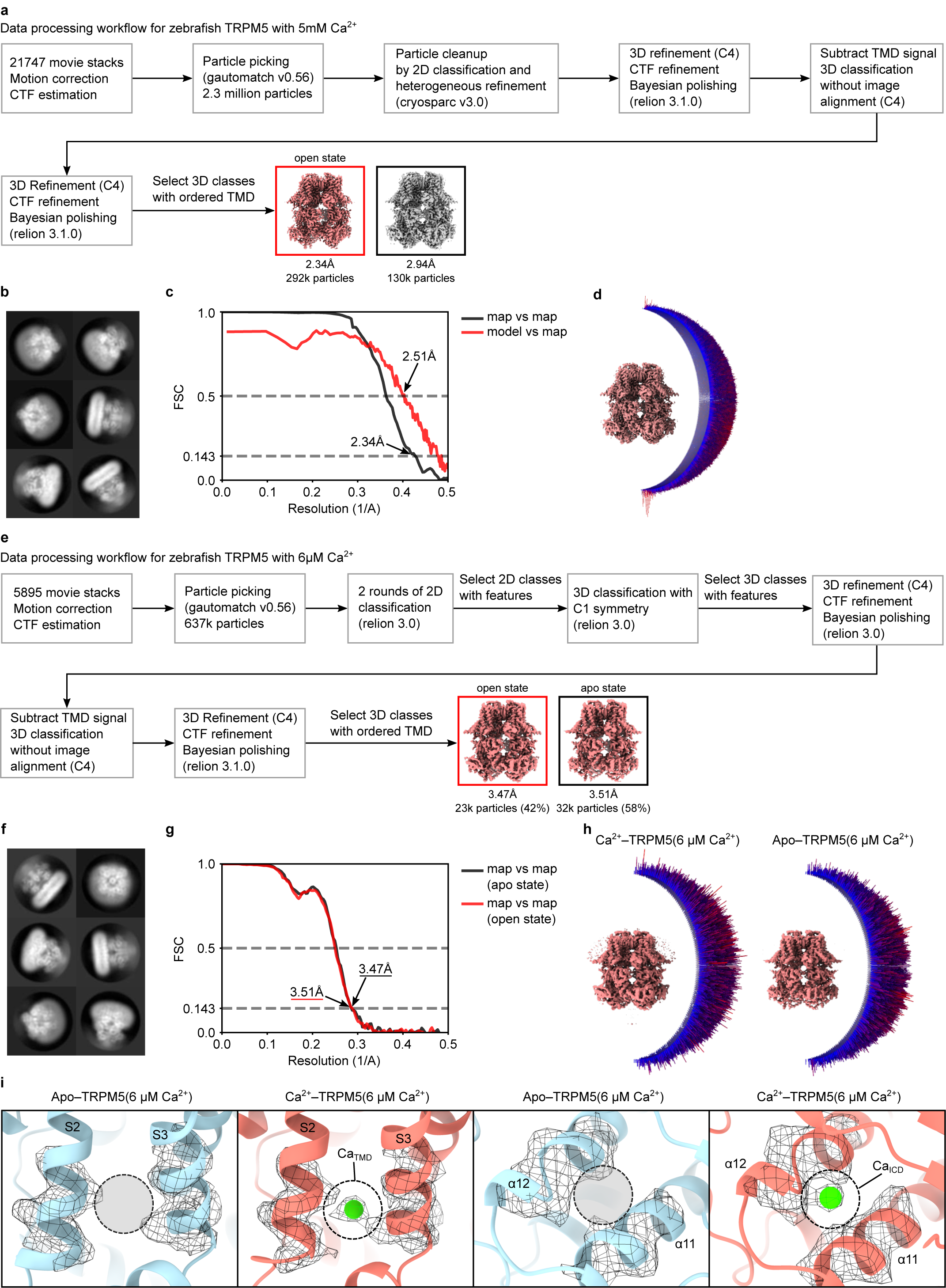
Ca^2+^–TRPM5 in GDN detergent. **a** and **e**, The data processing workflow for TRPM5 with 5mM Ca^2+^ dataset (**a**) and TRPM5 with 6 μM Ca^2+^ dataset (**e**). For the 5mM Ca^2+^ dataset, we obtained two conformations with ordered TMD after focused classification. While the Ca_TMD_ and Ca_ICD_ are fully occupied in both classes, there is subtle differences in the TMD. We focus on the conformation with the highest nominal resolution for model building and the discussion in the manuscript. **b** and **f**, The representative 2D class average the 5mM Ca^2+^ dataset (**b**) and 6 μM Ca^2+^ dataset (**f**), respectively. **c** and **g**, The Fourier shell correlation (FSC) curves for the 5mM Ca^2+^ dataset (**c**) and 6 μM Ca^2+^ dataset (**g**). The cryo-EM map FSC is shown in black and the model vs. map cross-correlation is shown in red. The map resolution is determined by the gold-standard FSC at 0.143 criterion, whereas the model vs. map resolution is determined by a correlation threshold of 0.5. **d** and **h**, The angular distribution of particles that give rise to the cryo-EM map reconstruction for 5mM Ca^2+^ dataset (**d**) and 6 μM Ca^2+^ dataset (**h**). **i**, Close up view of the Ca_TMD_ and Ca_ICD_ of the 6 μM Ca^2+^ dataset. From left to right, Ca_TMD_ of apo–TRPM5(6 μM Ca^2+^), Ca_TMD_ of Ca^2+^–TRPM5(6 μM Ca^2+^), Ca_ICD_ of apo–TRPM5(6 μM Ca^2+^), and Ca_ICD_ of Ca^2+^–TRPM5(6 μM Ca^2+^). The cryo-EM densities are shown in mesh representation. The expected Ca^2+^ density is indicated by a circle.

**Extended Data Figure 5:**
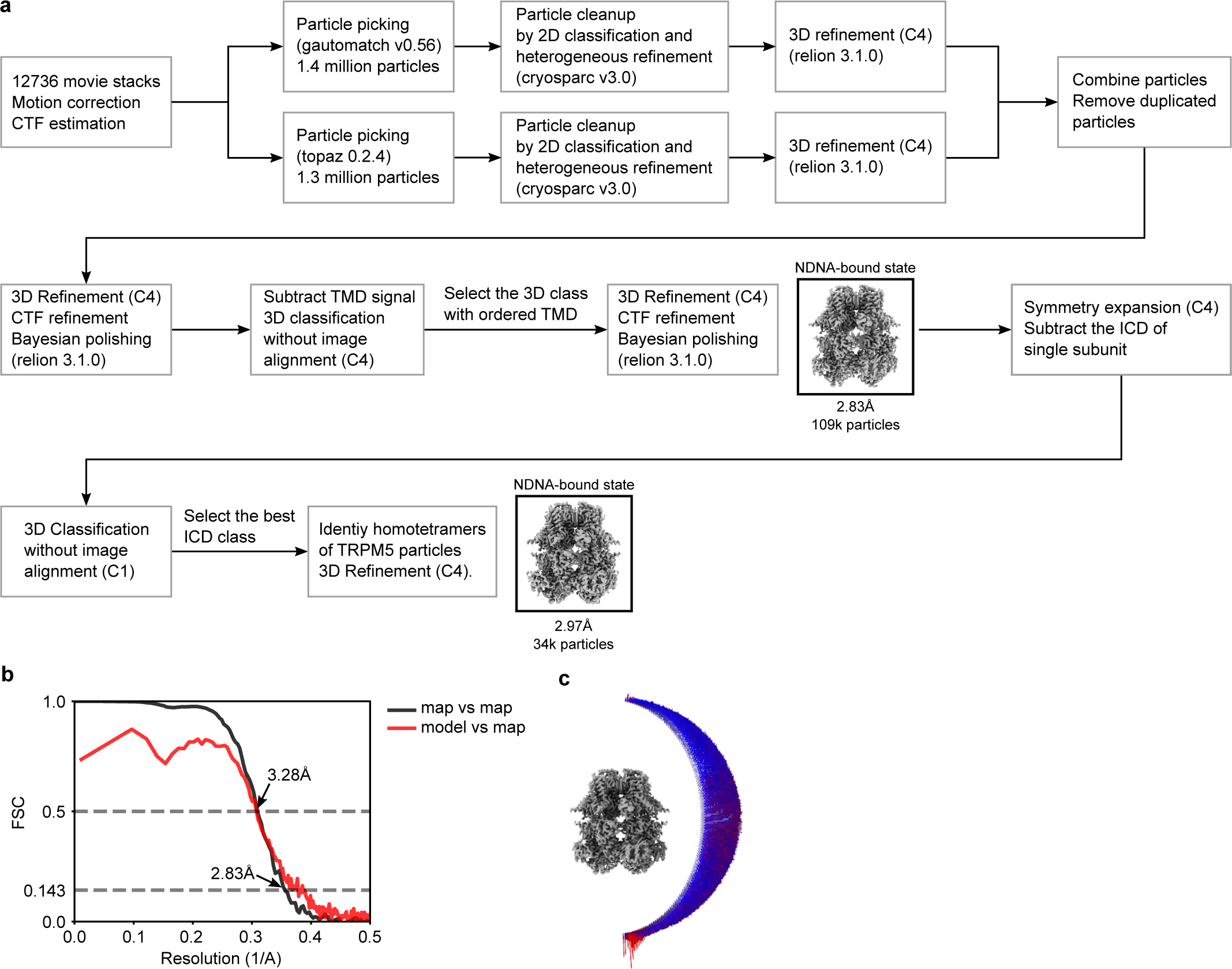
NDNA/Ca^2+^–TRPM5 in GDN detergent. **a**, The data processing workflow for NDNA/Ca^2+^–TRPM5 dataset. After TMD-focused classification and refinement, we found that the intracellular domain still contains major heterogeneity. To facililate model building, we analyzed the intracellular domain at the single subunit level through 3D classification. A set of single subunit particles with homogeneous ICD were converted back to the tetrameric TRPM5 particles and further refined to 2.97 Å resolution. Although the final map is of slightly worse nominal resolution (2.97 Å vs 2.83 Å), it facilitates the model building in the ICD. **b**, The Fourier shell correlation (FSC) curves for the NDNA/Ca^2+^–TRPM5. The cryo-EM map FSC is shown in black and the model vs. map cross-correlation is shown in red. The map resolution was determined by the gold-standard FSC at 0.143 criterion, whereas the model vs. map resolution was determined by a correlation threshold of 0.5. **d**, The angular distribution of particles that gave rise to the NDNA/Ca^2+^–TRPM5 cryo-EM map reconstruction.

**Extended Data Figure 6:**
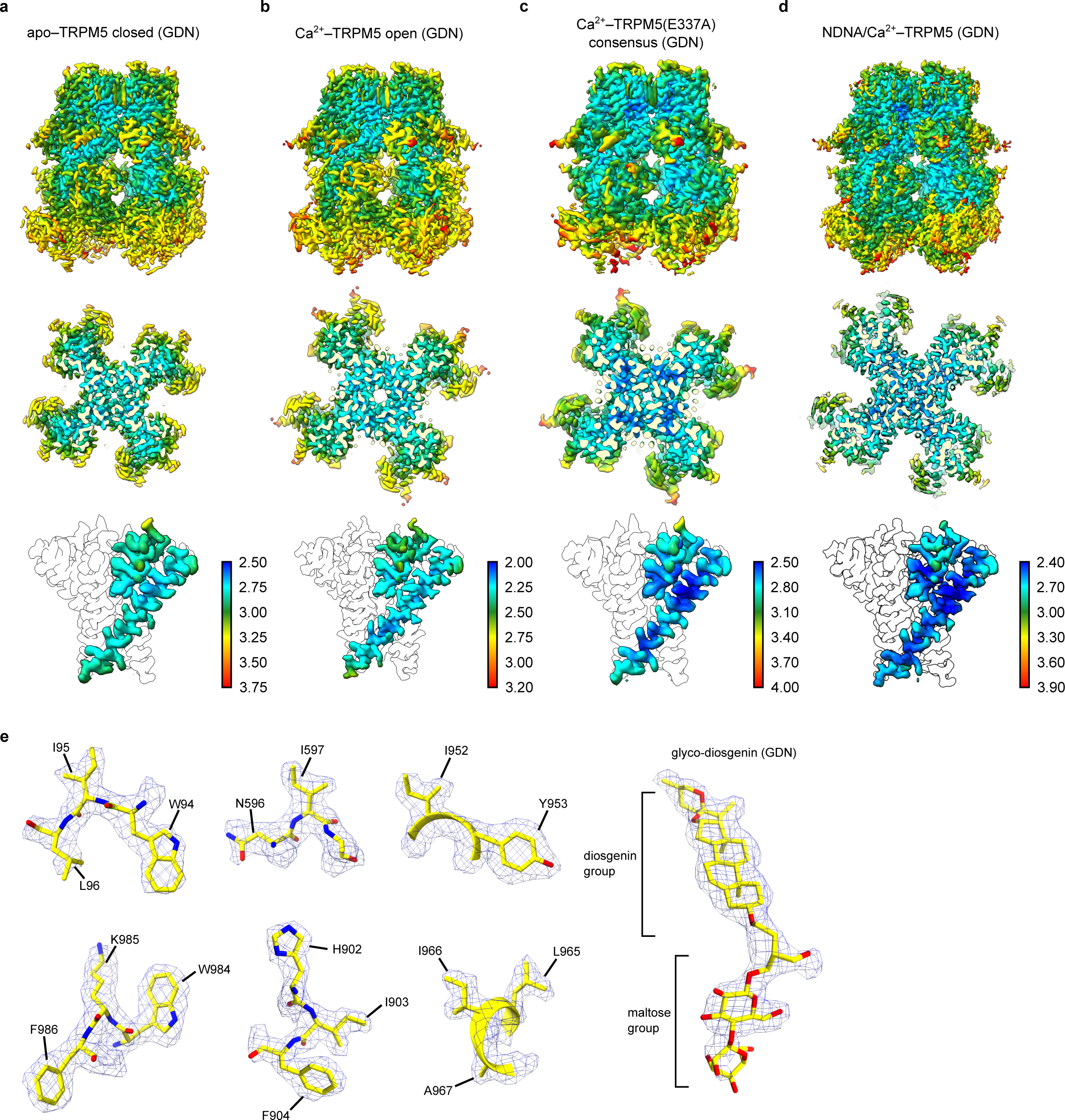
Local resolution estimation of TRPM5 structures and representative densities. **a-d**, The local resolution estimation for apo–TRPM5(GDN) (**a**), Ca^2+^– TRPM5(GDN) (**b**), Ca^2+^–TRPM5(E337A)(GDN) consensus (**c**), NDNA/Ca^2+^–TRPM5(GDN) (**d**). For each map, a side view, a top-down view of the TMD from the extracellular side, and a focused side view of the S6 and pore helix are shown. The color bar unit is in Ångstroms. **e**, Representative densities from Ca^2+^–TRPM5(GDN) map. For the GDN density, one maltose group of the molecule is not resolved in the cryo-EM density map.

**Extended Data Figure 7:**
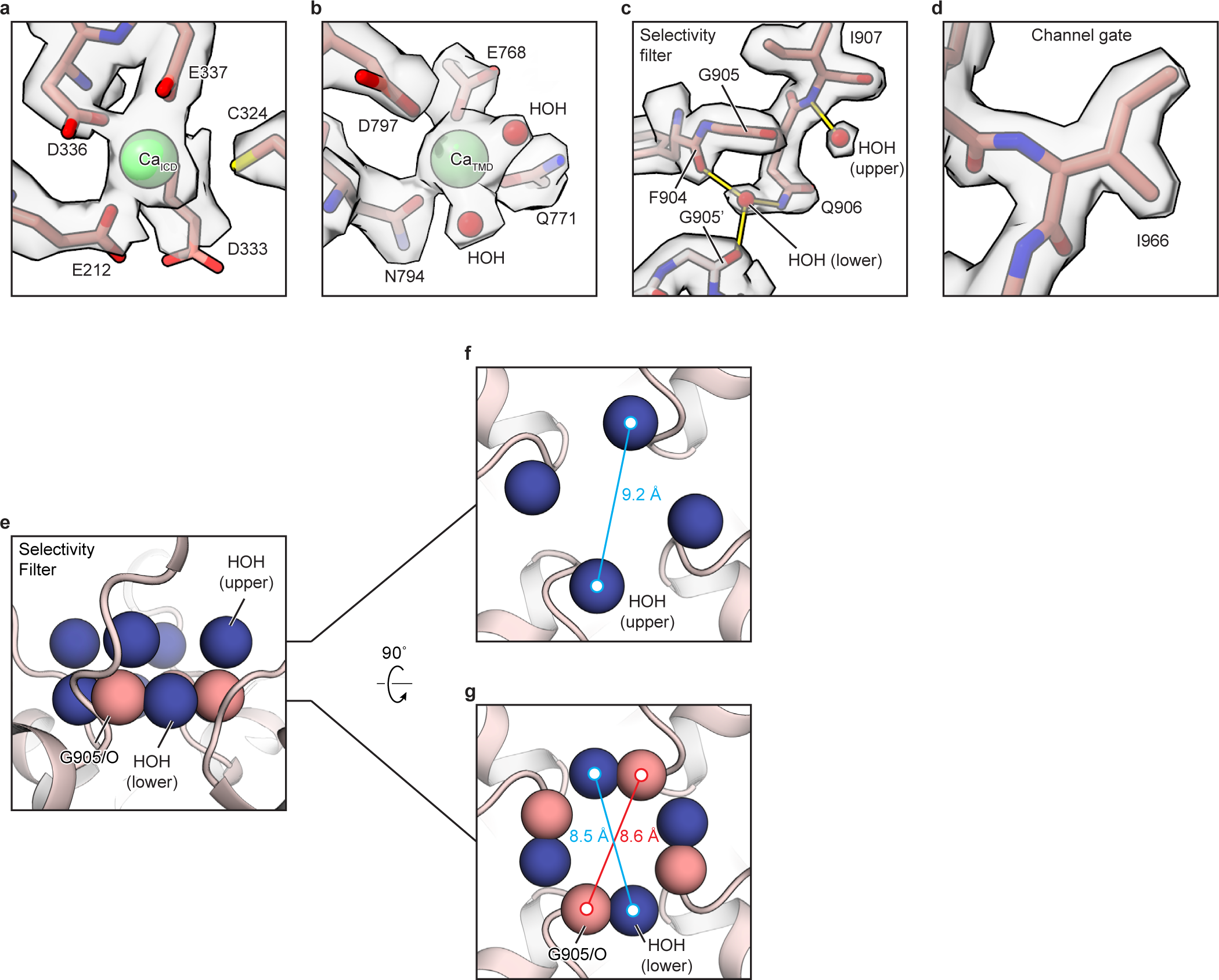
The gate and the selectivity filter of TRPM5. **a**, Cryo-EM densities of the Ca_ICD_ site, contoured at 0.018. **b**, Cryo-EM densities of the Ca_TMD_ site, contoured at 0.022. **c**, Cryo-EM densities of the water molecule and residues in the selectivity filter, contoured at 0.023. Hydrogen bonds are shown as solid yellow lines. The “lower” water molecule is surrounded by the sidechain of Q906 and the backbone oxygen atoms of F904 and G905, forming three hydrogen bonds. d, Cryo-EM densities of I966, which forms the channel gate, contoured at 0.03. **e**, The selectivity filter formed by two layers of ordered water molecules (blue spheres) and backbone oxygen atoms (pink spheres) of G905. **f** and **g**, The two hydration layers the selectivity filter viewed from the extracellular side. Upper layer in (**f**) and lower layer in (**g**).

**Extended Data Figure 8:**
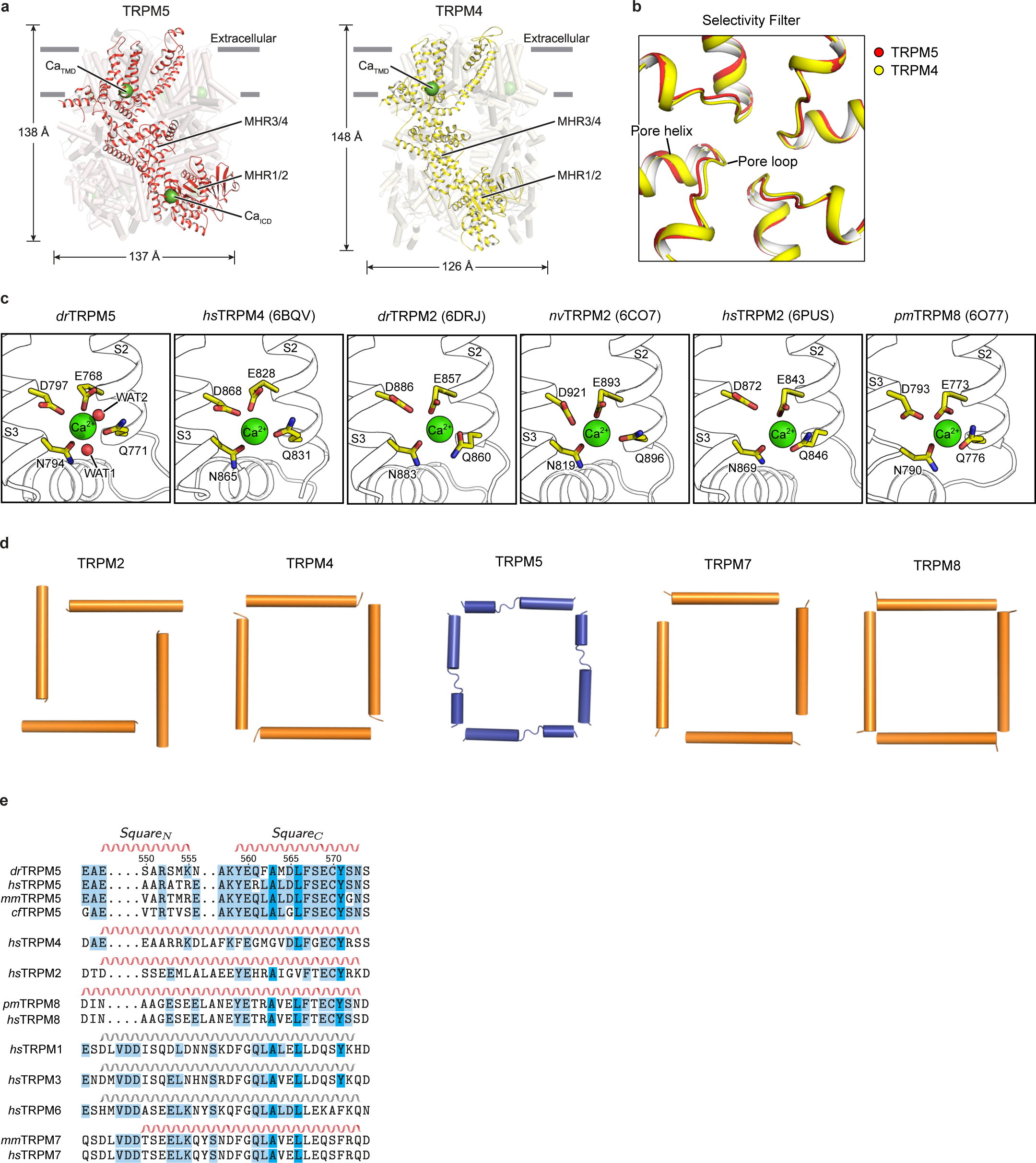
Comparison of TRPM5 with other TRPM channels. **a**, A structural comparison between Ca^2+^–TRPM5 and TRPM4 (PDBID: 6BQV). A single subunit is in color and shown as a cartoon. The TRPM5 channel is more compact, but wider than, the TRPM4 channel. **b**, An overlap of the selectivity filter of Ca^2+^–TRPM5 (red) and TRPM4 (yellow). **c**, A comparison of the Ca_TMD_ site for the available TRPM members. From left to right, *dr*TRPM5, *hs*TRPM4 (6BQV), *dr*TRPM2 (6DRJ), *nv*TRPM2 (6CO7), *hs*TRPM2 (6PUS), and *pm*TRPM8 (6O77)^23, 24, 31, 37, 38^; Shown in parentheses are the PDBIDs. **d**, Comparison of the “square” helices in TRPM2 (6PUO), TRPM4 (6BQR), TRPM5, TRPM7 (5ZX5), TRPM8 (6O6A); Shown in parentheses are the PDBIDs. Only TRPM5 has a broken square helix. **e**, A sequence alignment of the square helix across different TRPM5 orthologs and TRPM family members. Red indicates that the α-helix is observed in structures. For TRPM1, TRPM3, and TRPM6, in which no structures are currently available, the helical annotation is based on the secondary structure prediction from PSIPRED server^40^.

**Extended Data Figure 9:**
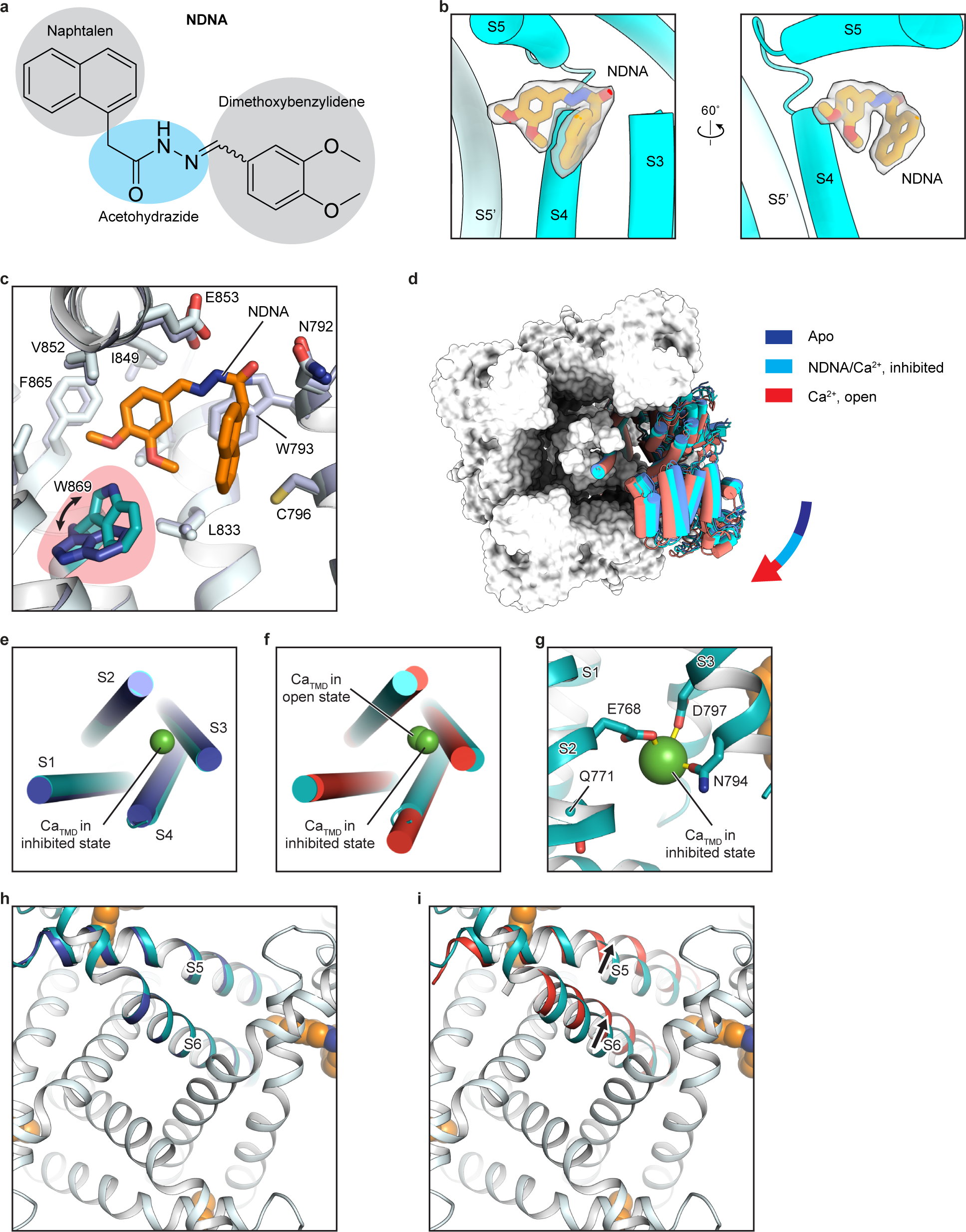
Comparison of NDNA/Ca^2+^–TRPM5 with apo–TRPM5 and Ca^2+^–TRPM5. **a**, the chemical structure of N’-(3,4-dimethoxybenzylidene)-2-(naphthalen-1-yl)acetohydrazide (NDNA). **b**, Two close-up views of the cryo-EM densities of NDNA molecule. The surrounding protein structural element is shown in cartoon representation. **c**, Comparison of the NDNA binding site between NDNA/Ca^2+^–TRPM5 and apo–TRPM5 structures. The W869 is flipped in the NDNA/Ca^2+^–TRPM5 structure (cyan) compared to that in apo–TRPM5 structure (blue). **d**, Overlay of NDNA/Ca^2+^–TRPM5 (cyan) with apo–TRPM5 (blue) and Ca^2+^–TRPM5 (red) structures view from the intracellular side. One subunit is shown in cartoon representation and the other three subunits are in surface representation. The ICD of NDNA/Ca^2+^–TRPM5 adopts an intermediate state compared to the apo–TRPM5 and Ca^2+^– TRPM5 structures. **e**, The superimposition of the S1-S4 domain between NDNA/Ca^2+^–TRPM5 (cyan) and apo–TRPM5 (blue). **f**, The superimposition of the S1-S4 domain between NDNA/Ca^2+^–TRPM5 (cyan) and Ca^2+^–TRPM5 (red). **g**, A close-up view of the Ca_TMD_ site in NDNA/Ca^2+^–TRPM5 structure. The Q771 moved away from Ca_TMD_. **h** and **i**, An overlay of the pore domain between NDNA/Ca^2+^–TRPM5 (cyan) with apo–TRPM5 (blue) (**h**) and Ca^2+^– TRPM5 (red) (**i**) structures viewed from the extracellular side.

**Extended Data Figure 10:**
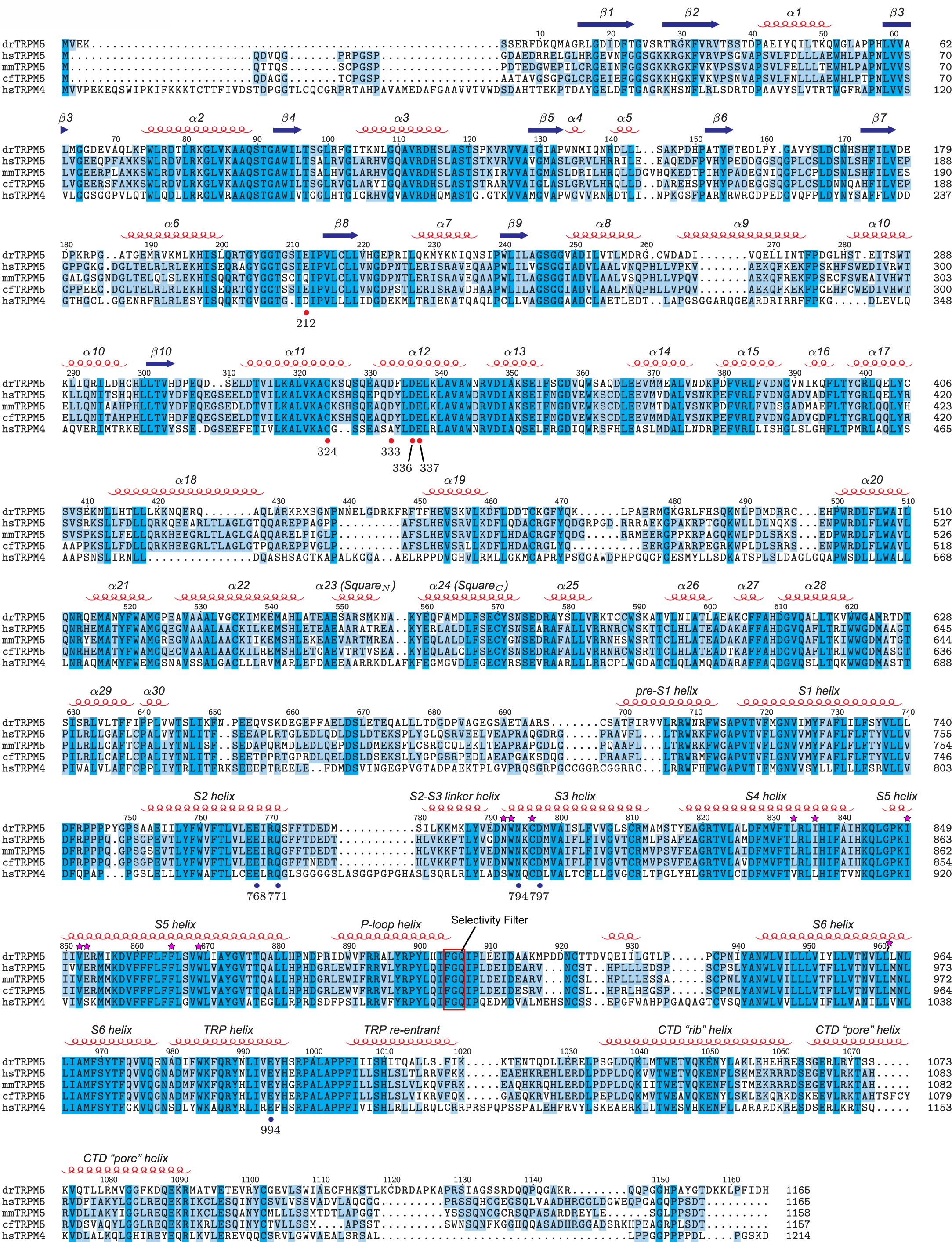
A sequence alignment of TRPM5 orthologs and *hs*TRPM4. Secondary structure elements are indicated at the top. Residues forming the Ca_TMD_ and Ca_ICD_ sites are indicated by blue and red dots, respectively. The selectivity filter is highlighted with a red frame. The NDNA interacting residues are marked with a magenta star.

**Extended Data Figure 11:**
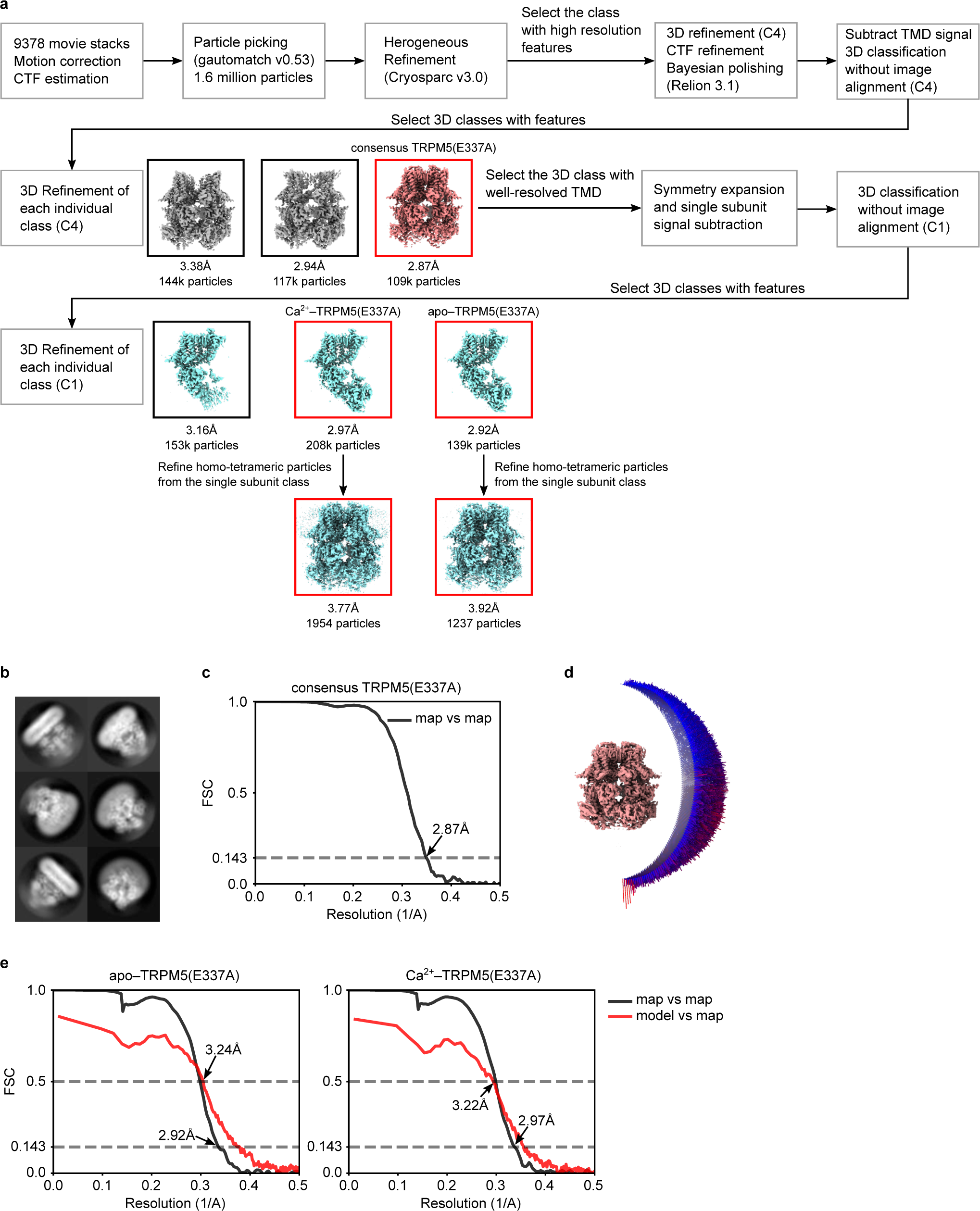
Ca^2+^–TRPM5(E337A) in GDN detergent. **a**, The data processing workflow for Ca^2+^–TRPM5(E337A) dataset. **b**, The representative 2D class average of Ca^2+^–TRPM5(E337A). **c**, The FSC curve for the consensus map of Ca^2+^–TRPM5(E337A). The map resolution was determined by the gold-standard FSC at 0.143 criterion. **d**, The angular distribution of particles that give rise to the consensus map of Ca^2+^–TRPM5(E337A). **e**, The FSC curve for apo–TRPM5(E337A) (left) and Ca^2+^–TRPM5(E337A) (right). For each panel, the cryo-EM map FSC curve is shown in black and the model vs. map corss-correlation is shown in red. The map resolution was determined by the gold-standard FSC at 0.143 criterion, whereas the model vs. map resolution was determined by a correlation threshold of 0.5.

**Extended Data Table 1:** Cryo-EM data collection, refinement, and validation statistics.

|  | Apo-TRPM5 | Ca<sup>2+</sup>-TRPM5 | Apo-TRPM5 (nanodisc) | Ca<sup>2+</sup>-TRPM5 (nanodisc) |
| --- | --- | --- | --- | --- |
| **Data collection and processing** |  |  |  |  |
| Magnification | 130,000 | 105,000 | 130,000 | 130,000 |
| Voltage (kV) | 300 | 300 | 300 | 300 |
| Electron exposure (e<sup>-</sup>/Å<sup>2</sup>) | 49.6 | 47 | 49.6 | 49.6 |
| Defocus range (μm) | -0.9 – -1.9 | -0.9 – -1.9 | -0.9 – -1.9 | -0.9 – -1.9 |
| Pixel size (Å) | 1.042 | 0.826 | 1.042 | 1.042 |
| Symmetry imposed | C4 | C4 | C4 | C4 |
| Initial particle images (no.) | 750k | 2,300k | 203k | 794k |
| Final particle images (no.) | 84k | 292k | 13k | 44k |
| Map resolution (Å) | 2.84 | 2.34 | 3.60 | 3.06 |
| FSC threshold | 0.143 | 0.143 | 0.143 | 0.143 |
| Map resolution range (Å) | 2.95 – 246.2 | 2.34 – 246.2 | 3.60 – 246.2 | 3.06 – 246.2 |
| **Refinement** |  |  |  |  |
| Initial model used (PDB code) | *De novo* | *De novo* |  |  |
| Model resolution (Å) | 3.14 | 2.51 |  |  |
| FSC threshold | 0.5 | 0.5 |  |  |
| Map sharpening *B* factor (Å<sup>2</sup>) | -70.22 | -43.99 |  |  |
| Model composition |  |  |  |  |
| Non-hydrogen atoms | 30708 | 30684 |  |  |
| Protein residues | 3984 | 3984 |  |  |
| Ligands | 12 | 32 |  |  |
| R.m.s. deviations |  |  |  |  |
| Bond lengths (Å) | 0.186 | 0.19 |  |  |
| Bond angles (°) | 1.039 | 1.053 |  |  |
| Validation |  |  |  |  |
| MolProbity score | 1.51 | 1.61 |  |  |
| Clashscore | 6.32 | 6.23 |  |  |
| Poor rotamers (%) | 0.00 | 0.00 |  |  |
| Ramachandran plot |  |  |  |  |
| Favored (%) | 97.06 | 96.10 |  |  |
| Allowed (%) | 2.94 | 3.90 |  |  |
| Disallowed (%) | 0.00 | 0.00 |  |  |

|  | Ca<sup>2+</sup>–TRPM5(E337A)<br>consensus | Apo–TRPM5(E337A)<br>subunit | Ca<sup>2+</sup>–TRPM5(E337A)<br>subunit |
| --- | --- | --- | --- |
| **Data collection and processing** |  |  |  |
| Magnification | 105,000 | 105,000 | 105,000 |
| Voltage (kV) | 300 | 300 | 300 |
| Electron exposure (e<sup>-</sup>/Å<sup>2</sup>) | 47 | 47 | 47 |
| Defocus range (μm) | -0.9 – -1.9 | -0.9 – -1.9 | -0.9 – -1.9 |
| Pixel size (Å) | 0.826 | 0.826 | 0.826 |
| Symmetry imposed | C4 | C1 | C1 |
| Initial particle images (no.) | 1,600k |  |  |
| Final particle images (no.) | 72k | 208k | 139k |
| Map resolution (Å) | 2.92 | 2.92 | 2.97 |
| FSC threshold | 0.143 | 0.143 | 0.143 |
| Map resolution range (Å) | 2.92 – 246.2 | 2.92 – 246.2 | 2.97 – 246.2 |
| **Refinement** |  |  |  |
| Initial model used (PDB code) |  | *De novo* | *De novo* |
| Model resolution (Å) |  | 3.24 | 3.22 |
| FSC threshold |  | 0.5 | 0.5 |
| Map sharpening *B* factor (Å<sup>2</sup>) |  | -70.89 | -70.72 |
| Model composition |  |  |  |
| Non-hydrogen atoms |  | 30760 | 30780 |
| Protein residues |  | 3984 | 3984 |
| Ligands |  | 12 | 16 |
| R.m.s. deviations |  |  |  |
| Bond lengths (Å) |  | 0.189 | 0.190 |
| Bond angles (°) |  | 1.041 | 1.08 |
| Validation |  |  |  |
| MolProbity score |  | 1.72 | 1.85 |
| Clashscore |  | 9.33 | 11.6 |
| Poor rotamers (%) |  | 0.00 | 0.00 |
| Ramachandran plot |  |  |  |
| Favored (%) |  | 96.55 | 95.99 |
| Allowed (%) |  | 3.45 | 4.01 |
| Disallowed (%) |  | 0.00 | 0.00 |

|  | NDNA/Ca<sup>2+</sup>-TRPM5 | Apo-TRPM5(6μM Ca<sup>2+</sup>) | Ca<sup>2+</sup>-TRPM5(6μM Ca<sup>2+</sup>) |
| --- | --- | --- | --- |
| **Data collection and processing** |  |  |  |
| Magnification | 105,000 | 130,000 | 130,000 |
| Voltage (kV) | 300 | 300 | 300 |
| Electron exposure (e<sup>-</sup>/Å<sup>2</sup>) | 47 | 49.6 | 49.6 |
| Defocus range (μm) | -0.9 – -1.9 | -0.9 – -1.9 | -0.9 – -1.9 |
| Pixel size (Å) | 0.826 | 1.042 | 1.042 |
| Symmetry imposed | C4 | C4 | C4 |
| Initial particle images (no.) | 1,400k | 637k | 637k |
| Final particle images (no.) | 109k | 32k | 23k |
| Map resolution (Å) | 2.83 | 3.51 | 3.47 |
| FSC threshold | 0.143 | 0.143 | 0.143 |
| Map resolution range (Å) | 2.83 – 246.2 | 3.51 – 246.2 | 3.47 – 246.2 |
| **Refinement** |  |  |  |
| Initial model used (PDB code) | *De novo* |  |  |
| Model resolution (Å) | 3.28 |  |  |
| FSC threshold | 0.5 |  |  |
| Map sharpening *B* factor (Å<sup>2</sup>) | -75.36 | -117.99 | -104.77 |
| Model composition |  |  |  |
| Non-hydrogen atoms | 30804 |  |  |
| Protein residues | 3984 |  |  |
| Ligands | 24 |  |  |
| R.m.s. deviations |  |  |  |
| Bond lengths (Å) | 0.2855 |  |  |
| Bond angles (°) | 1.46 |  |  |
| Validation |  |  |  |
| MolProbity score | 1.45 |  |  |
| Clashscore | 6.90 |  |  |
| Poor rotamers (%) | 0.00 |  |  |
| Ramachandran plot |  |  |  |
| Favored (%) | 97.69 |  |  |
| Allowed (%) | 2.31 |  |  |
| Disallowed (%) | 0.00 |  |  |

**Supplementary Figure 1: Tail current analysis of TRPM5 current**. **a-e**, Tail currents (for inside-out patch clamp experiments performed in Fig 1-2, Extended Data Figure 1) were plotted as a function of clamp voltage for *dr*TRPM5(WT), *dr*TRPM5(E337A), *dr*TRPM5(C324A), *dr*TRPM5(D333A), *dr*TRPM5(E212A) and *dr*TRPM5(D336A). Tail current amplitudes were measured at a clamp of -140 mV following activation voltages from -200 mV to +200 mV. For normalization (bottom row), a clamp of +200 mV was chosen. The number of cells used for analysis were: *dr*TRPM5(WT): 1 µM Ca^2+^ [11], 30 µM [5], 100 µM [3], 1000 µM [3], *dr*TRPM5(E337A): 1 µM Ca^2+^ [3], 30 µM [3], 100 µM [5], 1000 µM [4], *dr*TRPM5(C324A): 1 µM Ca^2+^ [5], 30 µM [5], 100 µM [5], 1000 µM [4], *dr*TRPM5(D333A): 1 µM Ca^2+^ [5], 30 µM [4], 100 µM [4], 1000 µM [3], *dr*TRPM5(E212A): 1 µM Ca^2+^ [4], 30 µM [3], 100 µM [4], 1000 µM [2] and *dr*TRPM5(D336A): 1 µM Ca^2+^ [3], 30 µM [3], 100 µM [3], 1000 µM [3].

**Supplementary Figure 2: Apo–TRPM5 and Ca^2+^–TRPM5 in nanodiscs. a** and **b**, The data processing workflow for apo-TRPM5(nanodisc) and Ca^2+^–TRPM5(nanodisc), respectively. **c** and **d**, the representative 2D class averages of apo–TRPM5(nanodisc) and Ca^2+^– TRPM5(nanodisc), respectively. **e**, The FSC curves for the apo–TRPM5(nanodisc) (black) and Ca^2+^–TRPM5(nanodisc) (red). The map resolution was determined by the gold-standard FSC at 0.143 criterion. **f**, The angular distribution of apo–TRPM5(nanodisc) and Ca^2+^–TRPM5(nanodisc) particles that give rise to the cryo-EM map reconstructions. **g**, The local map correlations between apo–TRPM5(nanodisc) vs. apo–TRPM5(GDN) and between Ca^2+^–TRPM5(nanodisc) vs. Ca^2+^–TRPM5(GDN). The color bar represents the correlation coefficient.

**Supplementary Figure 3: The raw gel images.** The raw SDS gel image to produce Extended Data Fig. 2c.

**Supplementary Figure 4: Synthesis of NDNA. a**, NDNA is synthesized by a two-step chemical reaction. **b**, The 400mHz H^1^ NMR spectrum of the intermediate compound 2 (upper panel) and NDNA (lower panel).

## Notes

### Competing Interest Statement

The authors have declared no competing interest.

