## Supplemental Figures for "Structures of TRPM5 channel elucidate mechanism of activation and inhibition"

#### Supplementary Figure 1

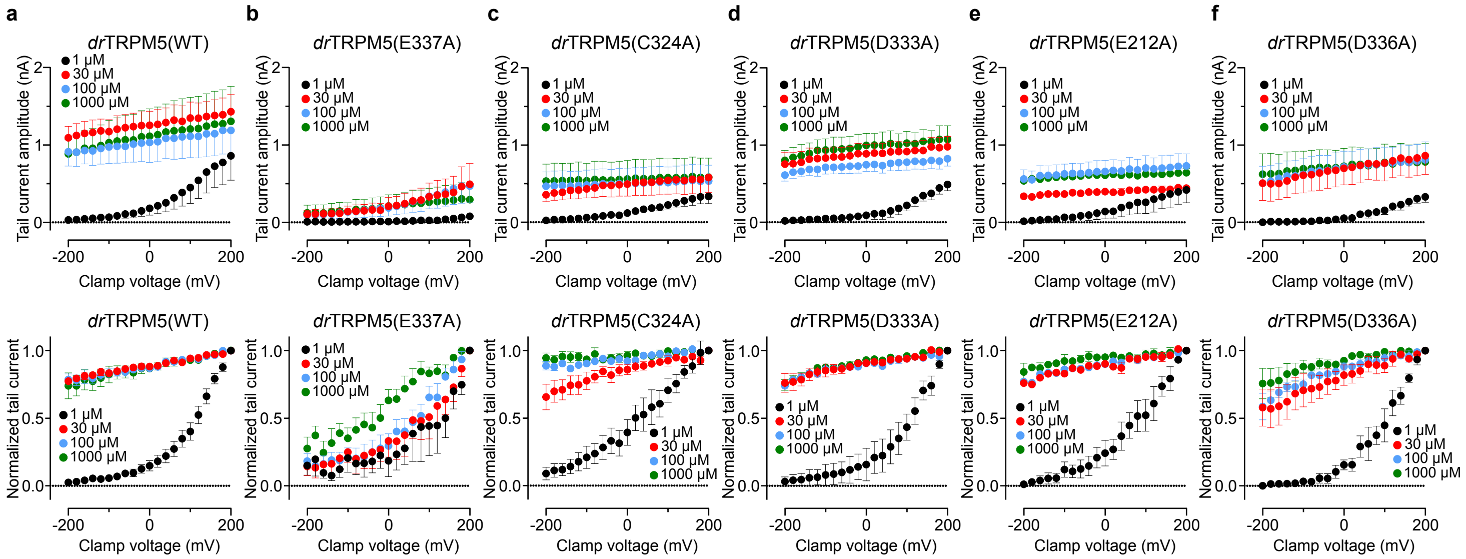

### Supplementary Figure 2

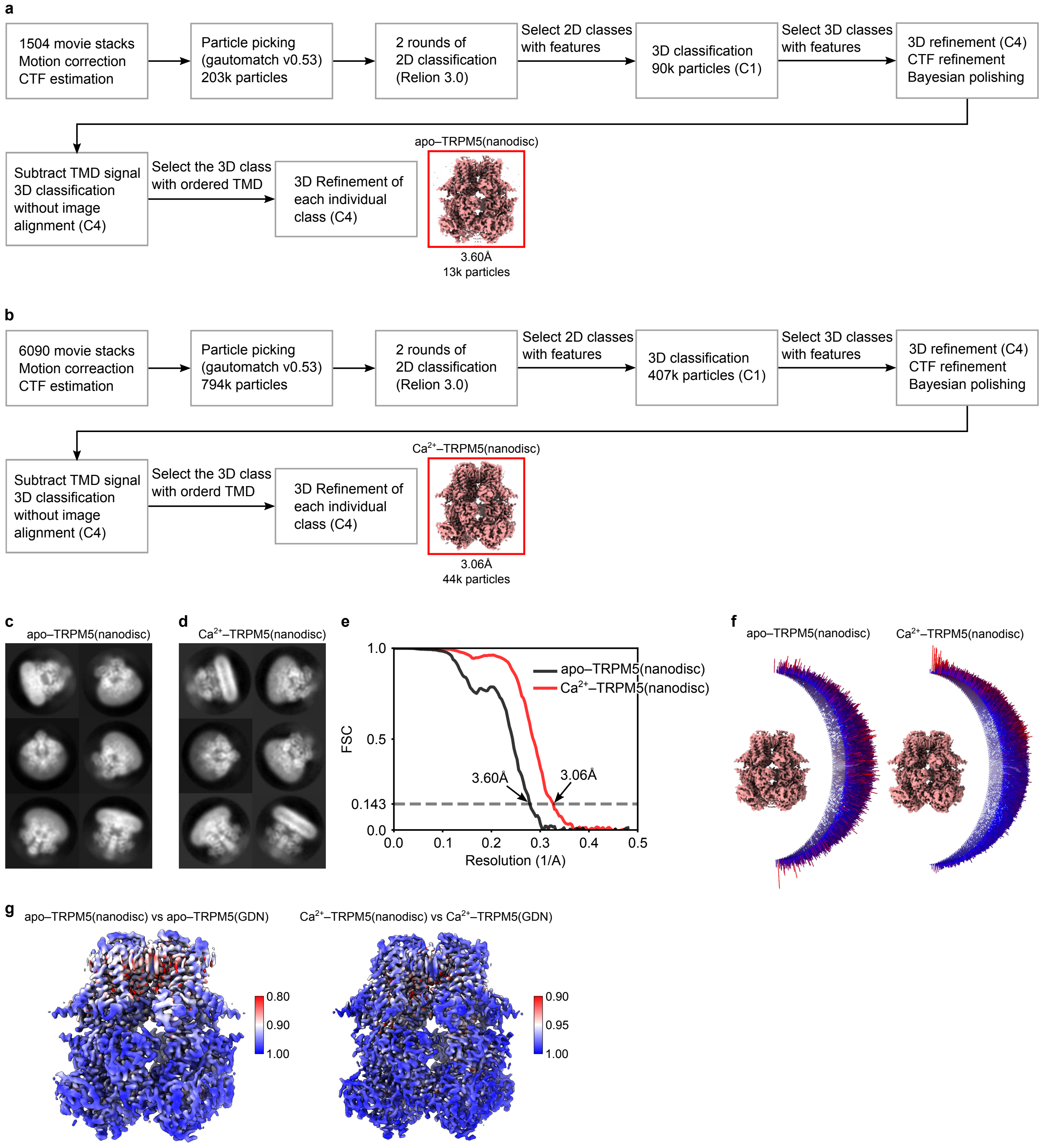

#### Supplementary Figure 3

Raw image for Extended Data Fig. 2c

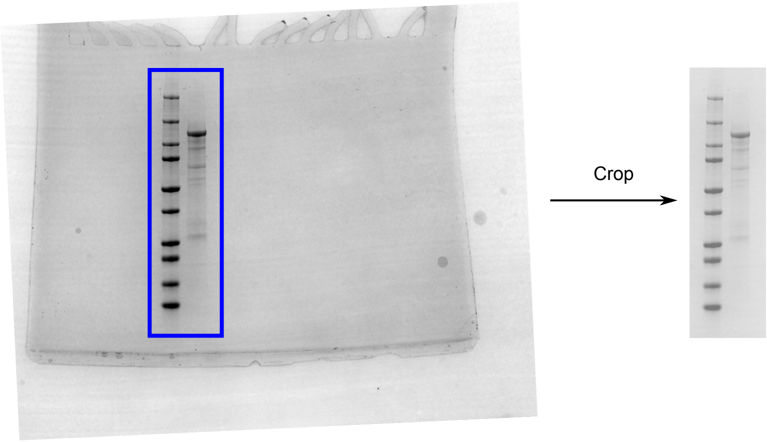

#### Supplementary Figure 4

**A**

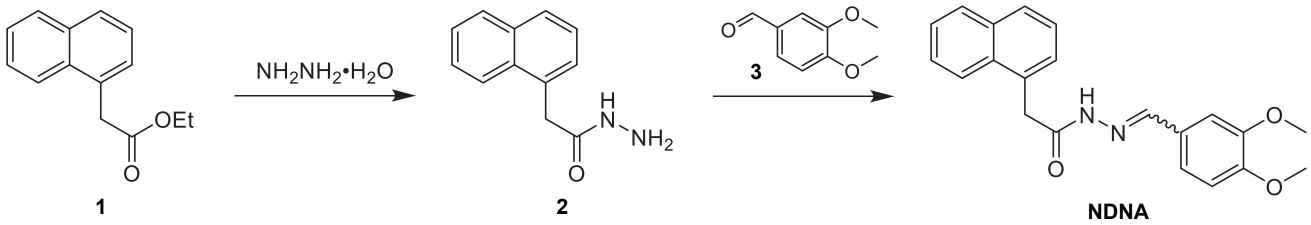


**B**

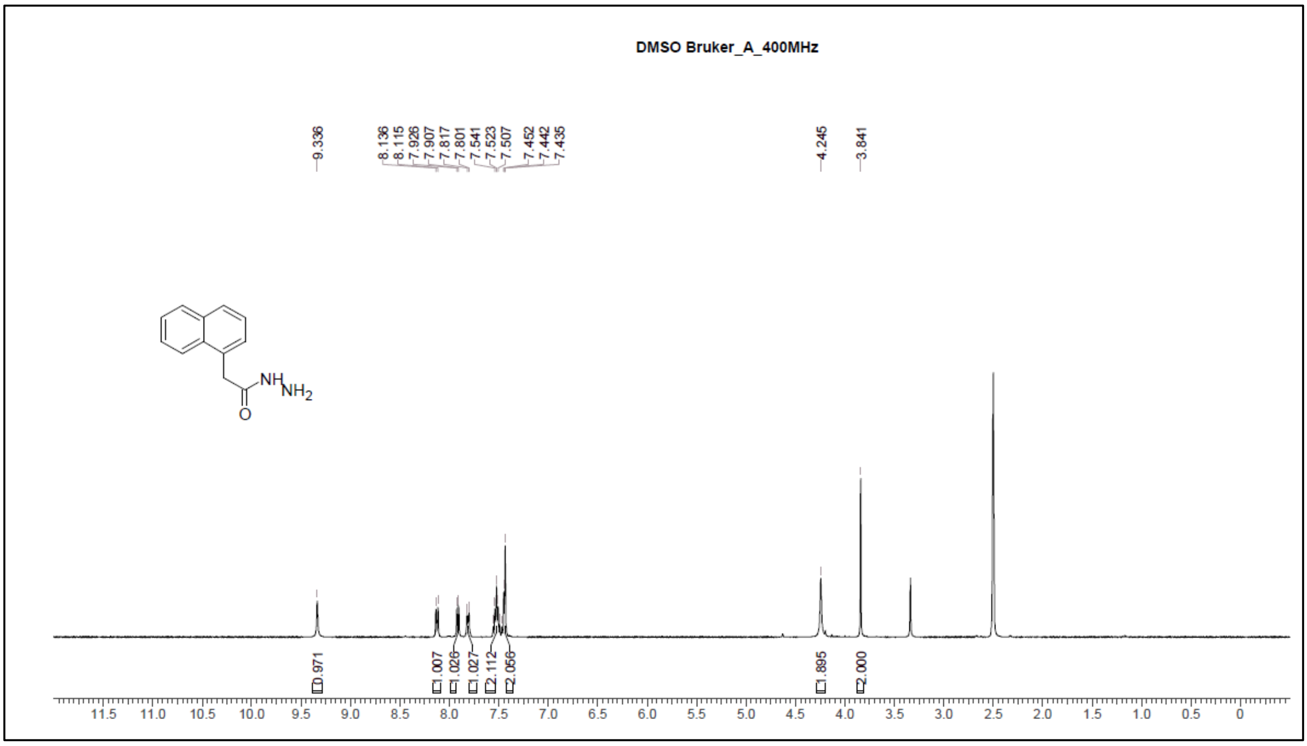


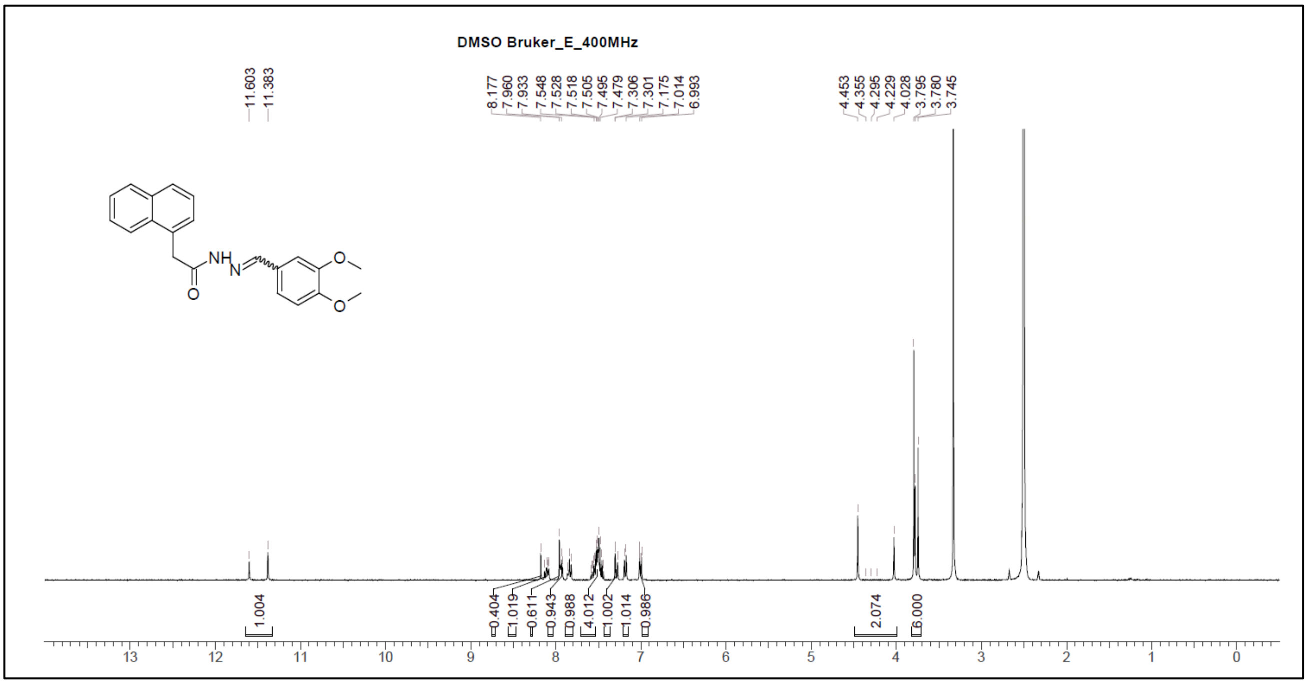
